# Ecophysiological and genomic approaches to cyanobacterial hardening for soil restoration

**DOI:** 10.1101/2023.09.07.556661

**Authors:** Roncero-Ramos Beatriz, Savaglia Valentina, Durieu Benoit, Van de Vreken Isabelle, Richel Aurore, Wilmotte Annick

## Abstract

Cyanobacteria inhabit extreme environments, including drylands, providing multiple benefits to the ecosystem. Soil degradation in warm drylands is increasing due to land-use intensification. Restoration methods adapted to the strong stress in drylands are being developed, i.e. cyanobacterial inoculation to recover biocrusts. For success, it is crucial to optimize the survival of inoculated cyanobacterial in field. One strategy is to harden them to be re-adapted to stressful conditions after laboratory culturing. Here, we analyzed the genome and ecophysiological response to osmotic, desiccation and UVR stresses of an Antarctic cyanobacterium, *Stenomitos frigidus* ULC029, closely related to other cyanobacteria from warm and cold dryland soils. Chlorophyll *a* concentrations show that preculturing ULC029 under moderate osmotic stress improved its survival during an assay of desiccation plus rehydration under UVR. Besides, its sequential exposition to these stress factors increased the production of exopolysaccharides, carotenoids and scytonemin. Desiccation, but not osmotic stress, increased the concentrations of the osmoprotectants, trehalose and sucrose. However, osmotic stress might induce the production of other osmoprotectants, for which the complete pathways were found in the ULC029 genome. In total, 140 genes known to be involved in stress resistance were annotated and could potentially help ULC029 under stress. Here, we confirm that the sequential application of moderate osmotic stress and dehydration, could improve cyanobacterial hardening for soil restoration, by inducing several resistance mechanisms. We provide a high-quality genome of ULC029 and a description of the main resistance mechanisms found (i.e. production of exopolysaccharides, osmoprotectants, chlorophyll and carotenoids; DNA repair; oxidative stress protection).

## Introduction

Cyanobacteria are extremophiles that can survive in different ecosystems, including some of the most extreme on Earth, such as the McMurdo Dry Valleys in Antarctica (Zhang et al. 2015) or the Atacama Desert in Chile (Patzelt et al. 2014). In terrestrial ecosystems, they are often part of communities, known as biological soil crusts or biocrusts, with other organisms (lichens, mosses, fungi, heterotrophic bacteria or microalgae) colonizing the first centimeter of soil surfaces (Weber et al. 2022). Biocrusts provide many ecosystem functions that improve soil fertility (Barger et al. 2016), stability (Belnap et al. 2007; Chamizo et al. 2017) and biodiversity (Maestre et al. 2011), among others. The cyanobacteria found in these ecosystems have mechanisms to withstand environmental stresses, such as high and continuous light intensity and UV radiation, desiccation, freezing or high salinity. They produce an exopolysaccharide matrix surrounding cells, which accumulates water, stabilizing the cell membrane during desiccation (Pereira et al. 2009). This EPS matrix also protects cell membranes from freezing-induced damage (Tamaru et al. 2005) and UVR (Chen et al. 2009), and accumulates nutrients (Mager and Thomas 2011), which is important to survive in oligotrophic environments. In order to prevent the production of reactive oxygen species (ROS) by UVA radiation, some cyanobacteria produce scytonemin, a yellow-brown pigment that is accumulated in the EPS matrix and which synthesis pathway has mainly been investigated in *Nostoc punctiforme* ATCC 29133 (Benett and Soule et al. 2022; Soule et al. 2013; Klicki et al. 2018). This pigment is the major contributor to surface temperature increase in warm deserts (Couradeau et al. 2016). In addition, cyanobacteria produce a wide range of mycosporine-like amino acids (MAAs) that accumulate inside cells and prevent DNA damage by absorbing UVB radiation (Gao and García-Pichel 2011). In addition to UVR, cyanobacteria in extreme environments are also exposed to high light intensity (continuously during summer in polar regions), which can lead to photooxidative damages by ROS formation (Muramatsu and Hihara 2012). In order to avoid ROS formation, chlorophyll concentration may be decreased, so that less energy is absorbed (Ogawa et al. 2018) and carotenoids may increase, harvesting light and acting as quenchers (Llewellyn et al. 2020; Gao and García-Pichel 2011). Energy can also be thermally dissipated through non-photochemical quenching by the orange-carotenoid-protein (OCP) (Muramatsu and Hihara 2012). Moreover, some motile cyanobacteria avoid solar radiation by vertical migration within mats or soil (Bebout and García-Pichel 1995). Another important stress factor in hot and cold deserts is the salinity. The two major strategies to withstand the osmotic stress due to an increase of the external ions concentration, present in a highly saline environment or due to events of desiccation or freezing, are the salt-out strategy and the production/uptake of osmoprotectants (Kvíderová et al. 2019). The former consists of exporting outside cells the ions that have entered, such as the toxic Na^+^, to maintain homeostasis whereas the latter promotes the intracellular accumulation of osmoprotectants to balance the osmotic potential and stabilize molecules structures (Kirsch et al. 2019). The type of osmoprotectants, also called compatible solutes, that cyanobacteria may produce or uptake, is roughly correlated to their natural habitat and their salt tolerance range (Kirsch et al. 2019). They are mainly sucrose and trehalose in low salt tolerance strains, glucosylglycerol in medium salt tolerance ones and glycine betaine and glutamate betaine for high salt tolerance strains (Mackay et al. 1984). However, there are a few exceptions (Kirsch et al. 2019). If these protective strategies do not succeed, the strategies to revert cellular damages are activated. Cyanobacteria can repair DNA damage by several mechanisms, such as the homologous recombination (HR) or the nucleotide excision repair (NER) pathways (Pathak et al. 2019), or refold proteins thanks to chaperones, such as GroEL/GroES or DnaK/J (Chatterjee et al. 2020). To revert ROS formation, they also produce oxygen-scavenging proteins, such as catalases or superoxide dismutases (Kvíderová et al. 2019).

Their ability to withstand extreme conditions makes them interesting organisms for different disciplines, such as astrobiology (Billi et al. 2022) or soil restoration in drylands (Rossi et al. 2017). The publications focused on dryland soil restoration by cyanobacterial inoculation have increased during the last decades (Antoninka et al. 2020) because of the increasing of degradation of dryland ecosystems due to climate change and anthropogenic impacts (Huang et al. 2016). In fact, the protection and restoration of terrestrial habitats have been included in the Sustainable Development Goals of the UN for 2030 (UN, 2015). Because cyanobacteria are the first colonizers of soils and thus participate in the first steps of biocrusts succession, soil restoration aims to induce the formation of biocrusts on degraded soils by inoculating different cyanobacteria (Weber et al. 2016). In laboratory conditions, the inoculated cyanobacteria can survive, colonize different types of degraded soils and induce biocrust formation (Adessi et al. 2021), thanks to their capacity to withstand the tested environmental stresses. However, when the restoration is tested in natural conditions in degraded dryland soils, the success rate decreases, even after hardening cyanobacterial cultures by, i.e., dehydration (Román et al. 2021a). Therefore, this study investigates the resistance mechanisms of a cyanobacterial strain to stresses encountered in their natural environment, with the aim to develop a hardening method that would succeed in their survival after inoculation.

For this goal, we followed a physiological and genomic approach. The selected strain, ULC029, had been isolated in one of the most extreme regions on Earth, continental Antarctica. This thin homocytous filamentous cyanobacterium was identified as *Leptolyngbya frigida* ANT.LH52B.3 at the time of isolation (Taton et al. 2006) but later renamed as *Stenomitos frigidus*, which belongs to a genus commonly found in dryland biocrusts (Patzelt et al. 2014; Pushkareva et al. 2015; Williams et al. 2016; Muñoz-Martín et al. 2019). Besides, this strain shows a high 16S rRNA gene similarity (99.4%; Shalygin et al. 2020) with several *Leptolyngbya* strains isolated in warm semiarid biocrusts in SE Spain (Roncero-Ramos et al. 2019). Moreover, it has been shown that the relative abundance of these strains increases in incipient biocrusts with ecosystem degradation (Roncero-Ramos et al. 2020). Also, other thin filamentous strains identified as *Leptolyngbya* have already shown promising results in inducing biocrusts after soil inoculation (i.e. Mugnai et al. 2018). On the other hand, authors studying the impact of stress have shown that the exposition to individual or combined stress factors may produce a different response (Rai et al. 2013; Joshi et al. 2017). Some stress factors might activate mechanisms usually associated with another one. For example, osmotic stress (Dillon et al. 2002) or desiccation (Fleming and Castenholz 2007) led to the production of the UV-screening pigment scytonemin. Therefore, we hypothesize that culturing cyanobacteria under multiple stress factors would improve their performance after inoculation in the field. To test our hypothesis, two ecophysiological assays were performed with cultures of ULC029 grown in the laboratory. First, we analyzed the influence of increasing salinity on cyanobacterial growth, the efficiency of photosynthesis *(F_v_/F_m_)* and the pigment production (chlorophyll *a*, carotenoids and scytonemin). After characterizing the salinity tolerance of the strain, we analyzed the influence of a multistressing environment including osmotic, desiccation and UVR stresses. In this second assay, we also quantified the osmoprotectants (sucrose and trehalose) and the exopolysaccharides. Finally, to identify all the potential resistance mechanisms of the strain ULC029, we sequenced and assembled the genome after a hybrid sequencing, which reduces the error rate and gaps, and thus, increases genome assembly quality (Koren et al. 2012). We identified the presence or absence of a selection of 140 genes known to be involved in cyanobacterial resistance mechanisms to environmental stress.

## Materials and Methods

### Cyanobacterial strain and culture conditions

The unicyanobacterial strain *Stenomitos frigidus* ULC029 (synonym *Leptolyngbya frigida;* Suppl. Figure 1), from the BCCM/ULC Cyanobacterial Collection, was isolated from a benthic microbial mat in the lake 52 (76°23’ S, 69°24’ E) in the Larsemann Hills in the Prydz Bay region of East Antarctica in 1998 (Taton et al. 2006). It was identified as *L. frigida* by morphological analysis and sequencing of the 16S rRNA gene and the ITS region (Taton et al. 2006), but it was later transferred to *S. frigidus* (Shalygin et al. 2020). A Maximum Likelihood tree (Suppl. Figure 2) built with all 16S rRNA sequences showing more than 96.3% similarity to ULC029 shows its close relatedness (circa 90%) with other strains from extreme biotopes (Antarctica and semiarid and dry environments). Concerning ULC029, the climatic conditions in the Larsemann Hills are characterized by a maximum snow annual precipitation of 250 mm, strong katabatic winds and air temperatures that range from maximum values of 4 or even 10 °C in summer and mean monthly temperatures between −15 and −18 °C in winter (Sabbe et al. 2004). When the lakes partially melt in summer, the water temperature can increase up to 8 °C. The lake 52 where the mat sample was collected is oligotrophic and oligosaline (conductivity: 138 µS · cm^−1^). The pH was 6.7 and the alkalinity 0.1 meq · L^−1^ (Sabbe et al. 2004).

ULC029 has been cultured at 12 °C under a continuous irradiance of 6 µmol photons m^−2^ s^−1^ and shaking at 80 rpm (IKA KS 4000). The culture medium was BG11 (Rippka et al. 1979).

### Experiment 1: salinity thresholds of *S. frigidus* ULC029

To determine the salt tolerance of the strain, it was cultured in BG11 with increasing concentrations of NaCl: 0, 0.5, 0.2, 0.7, 1.8 and 2.5 M. A homogeneous starting inoculum was prepared by centrifuging the cyanobacterial biomass at 4500 rpm for 7 minutes. The supernatant was discarded and the pellet resuspended in BG11 to reach a proportion of 20:80 of ULC029 pellet (solid) in BG11 (liquid). Then, 6 different inocula were prepared by mixing 1 mL of the prepared suspension of 20% of ULC029 in BG11 and 1 mL of solutions with increasing concentrations of NaCl in sterilized milliQ H_2_O. The final concentration of all compounds in BG11 after mixing the suspension with the increasing concentrations of NaCl was half for all of them. 12-multiwell plates were used to carry out the experiment. Each well was filled with 2 mL of inoculum and four replicates were set up for each treatment and time point (0 hours, 2 hours, 21 days and 32 days; Figure 1). All multiwell plates were incubated at 12 °C under an irradiance of 6 µmol photons m^−2^ s^−1^. At each time point, 4 replicates of each treatment were used to measure the efficiency of the photosynthesis (*F_v_/F_m_*) and the concentration of three pigments: chlorophyll *a*, carotenoids and scytonemin. At the end of the experiment, the cultures were subjected to one of the two additional steps: either a UVR treatment or a recovery phase. The first treatment was carried out by placing the multiwell plates containing four replicates of each treatment at day 32 inside a dark box provided with UVA, UVB and PAR lamps. The total irradiance received by the cultures was: 10 and 2 W·m^−2^ of UVA and B respectively, and 11 µmol photons m^−2^ · s^−1^ of PAR. The plates were left in the box during 3 days and after that, the same measurements as before were performed. For the second treatment, the biomass of four replicates of each treatment was transferred at day 32 to new multiwell plates filled with 2 mL of BG11 and was allowed to recover for 33 days. Then, the same two measurements were performed.

**Figure 1:**
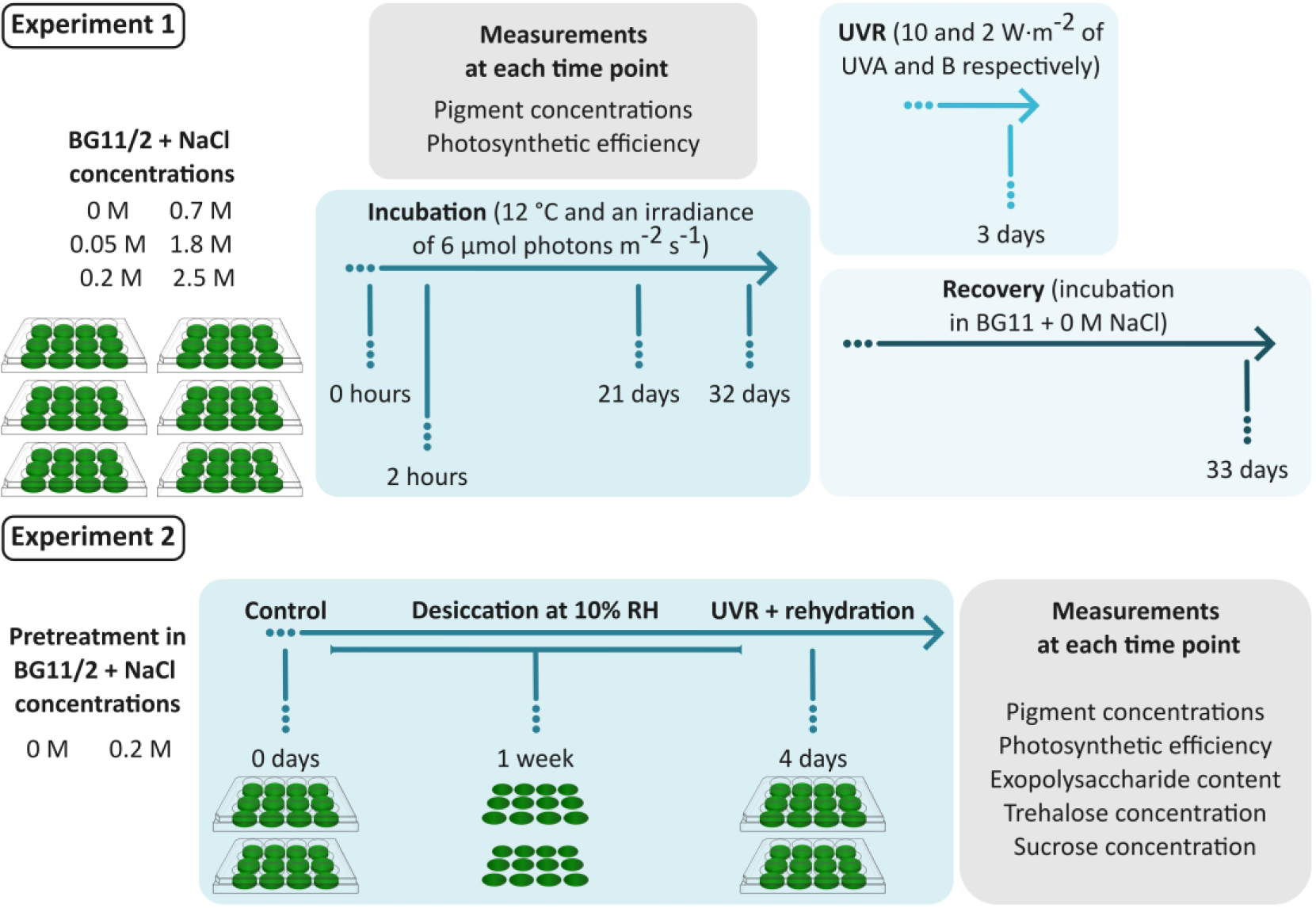
Experimental design of two ecophysiological assays: Experiment 1 and Experiment 2. During Experiment 1, ULC029 in BG11/2 and 6 concentrations of NaCl was incubated and monitored at four time points (0 hours, 2 hours, 21 days and 32 days). After 32 days, half of the replicates of each NaCl treatment were exposed to UVR during 3 days (4 replicates), and the other half incubated in BG11 + 0 NaCl during 33 days (Recovery phase). In Experiment 2, ULC029 cultures pre-treated in 0 or 0.2 M NaCl and transferred to multiwell plates (time 0 days, Control), were filtered and placed in a desiccation box during 1 week. Afterwards, filters were placed in multiwell plates and rehydrated with BG11 while exposed to UVR during 4 days.

### Experiment 2: Would a mild salinity stress improve the resistance of *S. frigidus* ULC029 to desiccation and UVR?

When ULC029 was cultivated in BG11 + 0.05 or 0.2 M NaCl, its growth and recovery appeared to be improved, as well as the resistance to UVR, following the data from Experiment 1. To better understand the potential effect of NaCl on the capacity of ULC029 to withstand stress, we carried on a second experiment with two media: BG11 (as a control) and BG11 + 0.2 M NaCl (the highest salt concentration allowing ULC029 to grow in Experiment 1; Figure 1). During a pre-adaptation phase, ULC029 was cultured in flasks with either culture medium (periodically refreshed) for more than 4 months. After that, as in Experiment 1, the biomass was centrifuged and resuspended in fresh medium to obtain two homogeneous inocula with the same quantity of biomass. 2 mL of each inocula were filtered on GF/F filters of Ø 25 mm (Whatman) for each replicate. Four replicates per treatment and measurement were set up. Three consecutive treatments were performed: the control which consisted of analyzing samples just after filtering; the desiccation treatment; and the rehydration + UVR treatment. The desiccation treatment consisted in transferring the filters into a chamber filled with 100 g of silica gel at the bottom to keep a HR of 10%, which allowed us to desiccate the filters in less than 1 day. Temperature and % RH inside the desiccation box were monitored with an iButton Hygrochron logger (Maxim Integrated, USA) every 5 minutes (Suppl. Figure 3). The filters were kept desiccated for 1 week. After that, the desiccation chamber was opened and four replicates were kept for each measurement. The rest of the filters were transferred to multi-well plates filled with 2 mL of BG11 medium to rehydrate them. Then, the plates were placed into the UV box with the same irradiances as in Experiment 1 and left inside during 4 days. After that, the last replicates were taken to perform all measurements. For this experiment, three additional variables were measured: the exopolysaccharides and the osmolytes (trehalose and sucrose) content.

### Culture monitoring and physiological measurements

In both experiments at each time point (including the initial inoculum), the culture growth was monitored by quantifying the chlorophyll *a* content as a proxy of photosynthetic biomass. It was extracted by mixing the biomass of each replicate with acetone 90% in a proportion of 1:5, vortexing and leaving it for 24 hours in darkness at 4 °C (following Castle et al. 2011 and Giraldo-Silva et al. 2020). After that, the samples were centrifuged (4500 rpm, 7 minutes at 4 °C) and the absorbance at 680 and 750 nm was measured with a UV-visible spectrophotometer (SPECORD 50, Analytik Jena). The quantity of two other pigments was also measured in the same extract at 490 (carotenoids) and 384 nm (scytonemin). The trichromatic equation was used to correct for the interferences of the other pigments (García-Pichel and Castenholz, 1991). The maximum photochemical <u>quantum yield</u> of photosystem II (*F_v_/F_m_*), considered as an indicator of photosynthesis efficiency and cellular stress, was measured with a PAM fluorometer (AquaPen AP-C, Photon Systems Instruments, CZ) after culture adaptation in the dark for 15 min.

Three additional variables were measured in Experiment 2, using 4 replicates per treatment and time point. The EPS were extracted to find out if their production was affected by the addition of NaCl to the medium, desiccation and/or the rehydration + UVR treatment. The extractions were divided in two categories: the loosely-bound EPSs (LB-EPSs) and the tightly-bound EPSs (TB-EPSs), following Rossi et al. (2018). The LB-EPSs were extracted by adding 3 mL of miliQ H_2_0, vortexing and incubating the filters for 15 minutes at 20 °C. After that, samples were centrifuged (4000 g, 20 min, 9 °C), the supernatant was transferred to a new tube, and the same process was repeated twice, ending up with a final extract of 9 mL, classified as LB-EPS fraction. After that, 3 mL of 0.1 M Na_2_EDTA was added to the filters, vortexed and incubated overnight at 20 °C. After centrifuging as described above, this step was repeated twice but decreasing the incubation time to 20 minutes. At the end, we got an extract of 9 mL, classified as the TB-EPS fraction. All the extractions were quantified by the phenolsulfuric acid method (Dubois et al. 1956), measuring the absorbance (488 nm) with a spectrophotometer (SPECORD 50, Analytik Jena). The total quantity of EPS was obtained by summing up both fractions. The other two variables measured in Experiment 2 were the quantification of two osmolytes: trehalose and sucrose. Filters were freeze-dried, eluted and measured with a Dionex System HPAED-PAD (High performance Anion Exchange Chromatography with Pulsed Amperometric Detection) equipped with a CarboPAC PA100 column of 4*250 mm. The eluents used were: A) HPLC-grade water + 100 mM NaOH, and B) HPLC-grade water + 100 mM NaOH + 600 mM of sodium acetate. The system was operated at 30 °C with a selected flow rate of 1 mL·min^−1^.

### DNA extraction and (meta)genome sequencing

A fresh culture of ULC029 in BG11 was used to extract genomic DNA using the GenElute Bacterial Genomic DNA Kit (Sigma-Aldrich) following the recommendations of the manufacturer. Three tubes with 1 mL of the culture were centrifuged (2000 rpm for 5 minutes), the supernatant discarded and washed again in BG11 several times to get rid of as many heterotrophic bacteria as possible, as it was not an axenic culture. At the end, we extracted the DNA from these three pellets. The integrity of the gDNA was controlled by electrophoresis and the DNA quality and concentration quantified with NanoVue (Biochrom). Finally, the three extractions were mixed to obtain enough DNA (> 5 µg) and sent to the sequencing platform (GIGA Genomics, University of Liege).

Both short and long reads of the gDNA were sequenced. Library preparation was performed using Nextera XT libraries (for Illumina sequencing) and Native barcoding genomic DNA kit (EXP-NBD114/LSK109) for Nanopore sequencing. Short reads (2×300 bp, 100x coverage) were sequenced by the Illumina Miseq v3 sequencing platform. Long reads sequences (coverage 100x) were obtained using a MinION Flow Cell R9.4.1 (Oxford Nanopore Technologies, UK).

### Bioinformatic analyses

As ULC029 culture was not axenic, its genomic dataset was treated as a metagenome and binned to obtain cyanobacterial specific contigs. The Illumina reads were filtered and trimmed using iu-filter-quality-minoche from Anvi’o (Eren et al. 2013), and their quality was checked with fastQC (Andrews et al. 2010). After controlling the Nanopore sequences quality with NanoPlot (v1.29.0; De Coster et al. 2018), NanoFilt (v2.6; De Coster et al. 2018) was used to filter out reads with a quality score lower than 10 and a length shorter than 500 bp. Long reads were assembled using Flye (metagenome mode; v2.9; Kolmogorov et al. 2020). Afterwards, filtered Illumina sequences were aligned to the assembled genome with bwa (v0.7.12; Li and Durbin 2009). The obtained sam files were converted to bam format and indexed using Samtools (v1.3.1; Danecek et al. 2021). Then, the assembled genome was polished with pilon (v1.23; Walker et al. 2014) using the indexed bam files as input. Binning was performed with CONCOCT (v1.1; Alneberg et al. 2014). Finally, we used GTDBtk (Parks et al., 2018) to determine the taxonomic placement of the bins and CheckM (v.1.1.6; Parks et al. 2014) to assess their completeness and contamination level. The assembly statistics were obtained with QUAST (default settings; v2.3; Gurevich et al. 2013). The genome was plotted using Proksee (Stothard and Wishart 2005) and manually edited with Inkscape v1.2. The genome assembly has been deposited at GenBank (Bioproject number: PRJNA1006388).

#### Genome annotation

A contig database was created using Anvi’o (Eren et al. 2015, 2021). Then, a first annotation was performed using a web version of Dfast (Tanizawa et al. 2016). Afterwards, genes in the database were annotated with functions from the NCBI’s Clusters of Orthologous Groups (COGs) using DIAMOND (fast mode; Buchfink et al. 2021) and with KEGG (Kyoto Encyclopedia of Genes and Genomes) identifiers using GhostKOALA (Kanehisa et al. 2016). Gene annotations and sequences are available in Suppl. Tables 1 and 2, respectively.

#### Identifying genes related with stress resistance

Genes known to be involved in resistance mechanisms against stress (Rajeev et al. 2013; Soule et al. 2013; Shimura et al. 2015; Al-Hosani et al. 2015; Kopf et al. 2015; Ferreira et al. 2016; Sinetova et al. 2016; Chrismas et al. 2016, 2018; Klicki et al. 2018; Shang et al. 2019; Pathak et al. 2019; Kvíderová et al. 2019; Pereira et al. 2009, 2019; Chatterjee et al. 2020; Mosca et al. 2021; Ye et al. 2021; Napoli et al. 2021) were searched in the generated contig database and classified in the following groups: (i) Production of the EPS matrix, (ii) Photoprotection, (iii) Oxidative stress protection, (iv) DNA repair and (v) Osmotic stress protection. When genes were not found in the database, they were searched manually using tblastn against the ULC029 assembled genome with an e-value threshold of 1e-5 and a score higher than 50 bits (following Pearson et al. 2013). The queries used can be found in the supplementary material (Suppl. Table 3). Alignments were validated by checking if the Pfam functional protein domains found with SMART (normal mode; Letunic et al. 2020) were the same as those from the query.

### Statistical analyses

In Experiment 1, results were analyzed to determine the effect of the increasing NaCl concentration on the pigments’ concentrations at different time points individually. Values of each time point were tested for normality and homoscedasticity with the Shapiro-Wilk (shapiro.test function, stats package) and Levene (function leveneTest, car package) tests, respectively. A two-way ANOVA (anova function, car package) followed by the Tukey post hoc test (TukeyHSD function, stats package) was applied when data followed a normal distribution and was homoscedastic, otherwise the Kruskal Wallis test (package conover.test) was applied. Moreover, the pigments contents after incubating the cultures under UVR from day 32 to 35 were analyzed by applying a one-way ANOVA (anova function, car package) and the Tukey post hoc test (TukeyHSD function, stats package) for chlorophyll *a* and carotenoid values independently.

Data obtained in Experiment 2 were also tested for normality and homocedasticity, as previously described. To determine the influence of the desiccation and rehydration+UVR treatments in cyanobacteria cultured in media with different NaCl concentrations, the following variables were analyzed: chlorophyll *a*, carotenoids, scytonemin, trehalose, sucrose and exopolysaccharides content. In all cases, we performed a two-way ANOVA (anova function, car package) followed by the Tukey posthoc test (package lsmeans).

Figures were plotted using the ggplot2 package, and manually edited using Inkscape v1.2. All analyses were performed with R 4.0.5 (Development Core Team 2017).

## Results

### Determining the salt tolerance of *S. frigidus* ULC029

*Stenomitos frigidus* ULC029 could grow and survive during 32 days in BG11 medium with a NaCl concentration up to 0.7 M (Figure 2a). After 32 days, the highest chlorophyll *a* concentration was found in medium with 0.2 M NaCl, followed by 0.05 M and 0 M. For NaCl concentrations of 1.8 and 2.5 M, the chlorophyll *a* concentration decreased with time and was null after one month. Moreover, there was no recovery as no chlorophyll *a* was found after transferring and incubating the cultures in BG11 without NaCl for another 33 days. Interestingly, the highest chlorophyll *a* values after recovery were found in cultures growing in 0.05 and 0.2 M NaCl, followed by 0 M NaCl (Figure 2a). However, although *S. frigidus* kept producing chlorophyll *a* in BG11 + 0.7 M NaCl during the recovery, it was significantly lower than with lower salt concentrations (Figure 2a). Therefore, 0.7 M NaCl is the maximum concentration that *S. frigidus* can withstand among the tested ones. In Figure 2b, the carotenoids concentration pattern is very similar to the one followed by chlorophyll *a*. In conclusion, the highest concentrations of chlorophyll *a* were obtained in 0.2, 0.05 and 0 M of NaCl (Figure 2b) after cultivation for 32 days, but also when a subsequent recovery was performed in 0 M NaCl for 33 days. Scytonemin was not detected in any treatments of Experiment 1.

**Figure 2:**
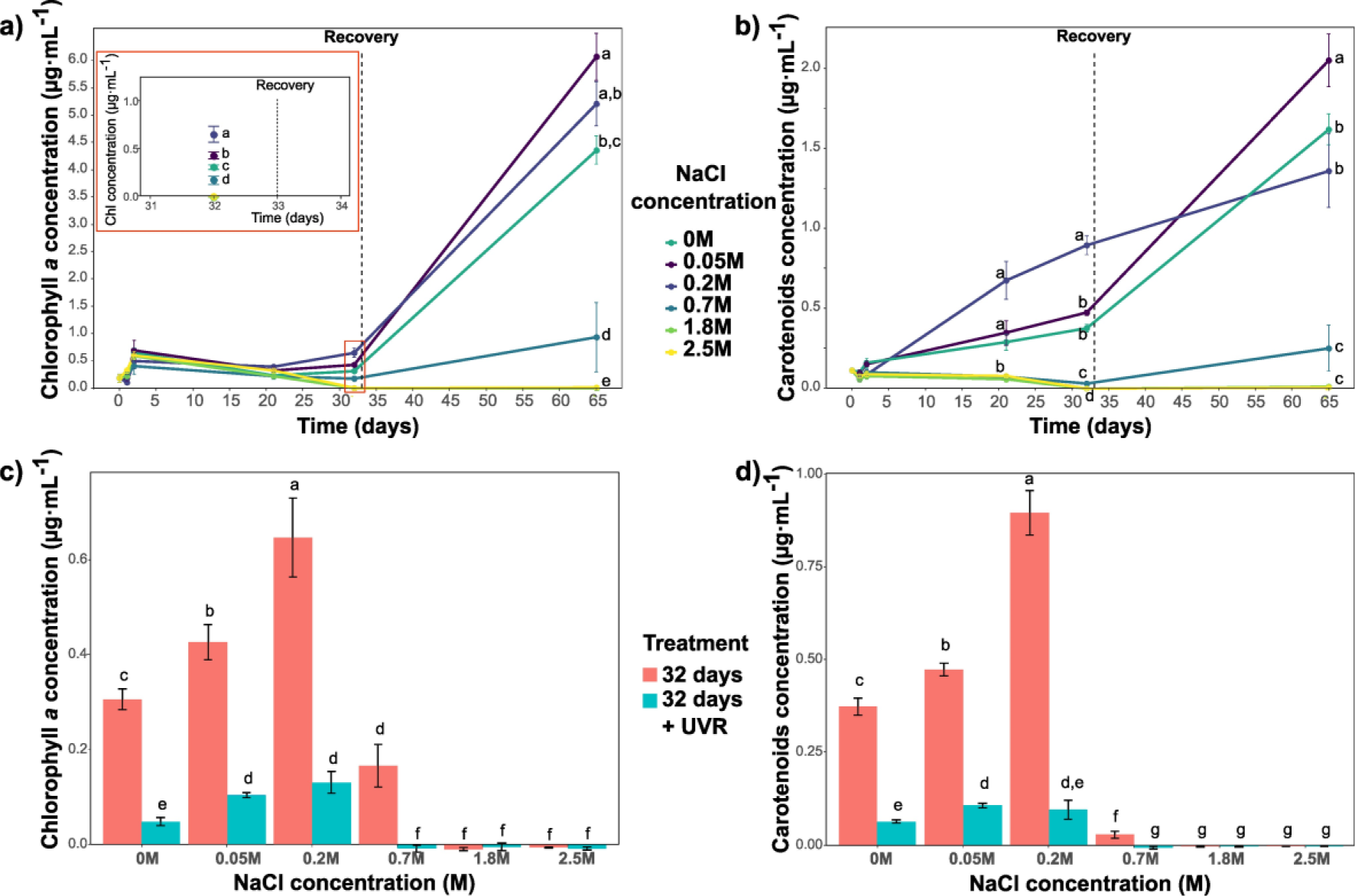
Pigments contents produced by *S. frigidus* ULC029 cultured at different concentrations of NaCl during 32 days of incubation and after subsequent recovery in BG11 medium for 33 days in Experiment 1: chlorophyll *a* (a) and carotenoids (b) concentrations during the 32 days of incubation; and chlorophyll *a* (c) and carotenoids (d) concentrations after 32 days compared with the same cultures exposed to three additional days of UVR. All data are means (n = 4). The error bars represent ±SE and letters indicate significant differences (p < 0.05) among treatments.

*F_v_/F_m_* values (∼ 0; Suppl. Figure 4) showed that *S. frigidus* was photosynthetically inactive in 0.7, 1.8 and 2.5 M NaCl during the growth phase. Only the strain cultured in 0.7 M NaCl could recover after 33 days in BG11 medium and became photosynthetically active again (*F_v_/F_m_* 0.3; Figure 2). Cultures in BG11 with 0.2 M NaCl were also photosynthetically inactive at day 32 (∼ 0; Suppl. Figure 4), but they could fully recover and reach the same values as at time 0 (∼ 0.2; Suppl. Figure 4). Finally, cultures with a low salinity stress (0.05 M NaCl) and the controls (0 M NaCl) were always photosynthetically active and both followed a similar pattern with the highest value at day 21 (0.3 and 0.4, respectively; Suppl. Figure 4).

When cultures were exposed during 3 days to UVR after 32 days of cultivation in media with different NaCl concentrations, both chlorophyll *a* and carotenoids concentrations were significantly lower compared to the non-exposed replicates (Figure 2c and d). However, cultures previously grown in media with 0 and 0.05 NaCl did not show stress and were photosynthetically active even after UVR exposure (Suppl. Figure 4). Also, cultures in 0.2 M NaCl showed higher *F_v_/F_m_* values, and thus were more photosynthetically efficient, after 3 days of UVR compared to the control which was measured before UVR exposure (Suppl. Figure 4).

### Ecophysiological response of *S. frigidus* ULC029 to multiple stress factors (osmotic stress pretreatment, desiccation and UVR)

First of all, contrary to the results obtained in Experiment 1, none of the variables measured showed a significant difference between the cultures cultivated in medium with 0 or 0.2 M NaCl in the absence of other stresses (Control treatment; Figure 3). However, significant differences were found in most of variables after desiccating the cultures for one week and the 4 days of rehydration and UVR exposure.

**Figure 3:**
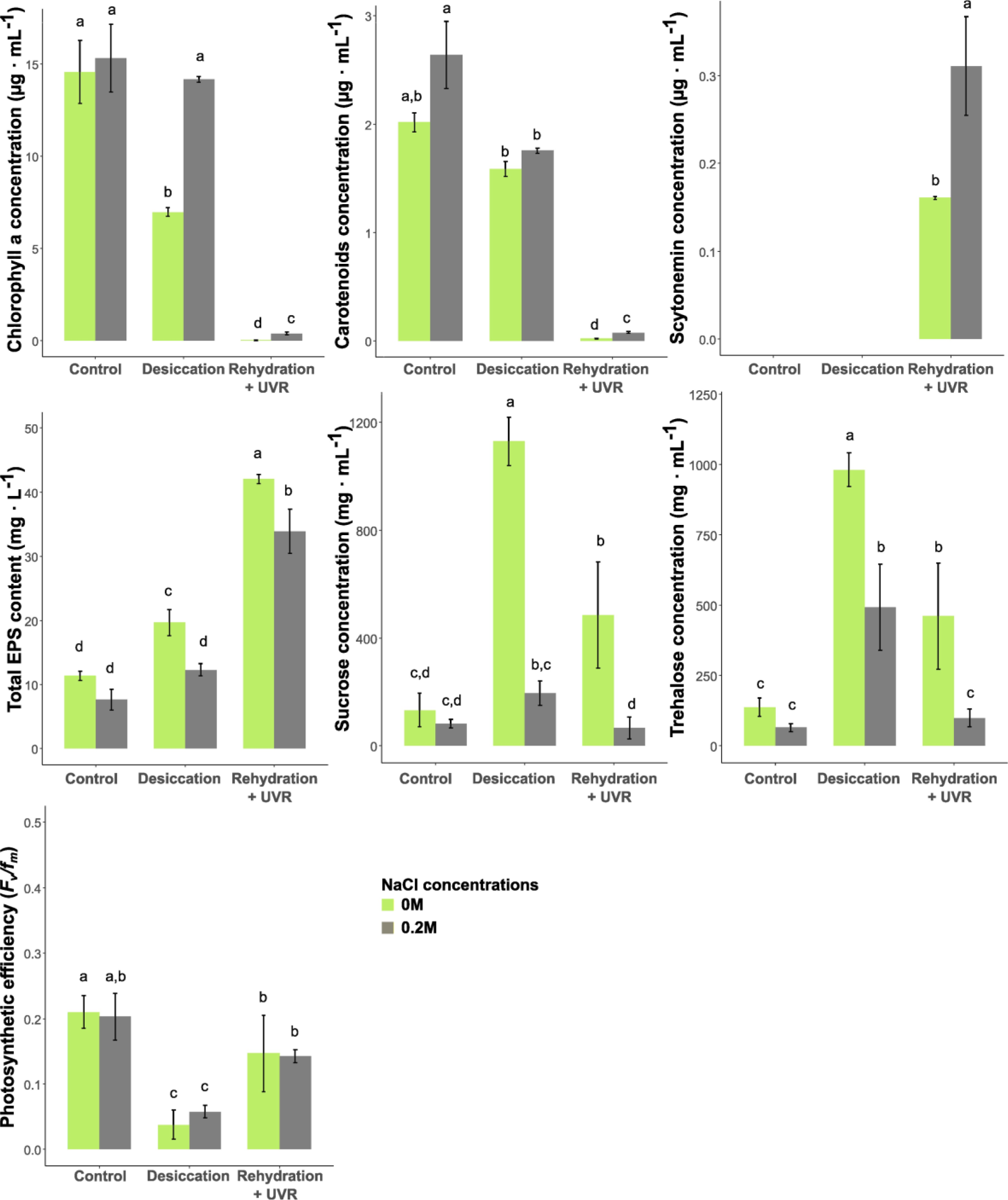
Influence of 1 week of desiccation and subsequent 4 days of rehydration combined with UVR exposure on *S. frigidus* ULC029 cultured in BG11 with 0 or 0.2 M NaCl compared to the control (initial values before desiccation) on different variables: chlorophyll *a*, carotenoids, scytonemin and total exopolysaccharides contents; sucrose and trehalose concentrations; and the photosynthetic efficiency (*F_v_/f_m_*). All data are means (n = 4). The error bars represent ±SE and letters indicate significant differences (p < 0.05) among treatments.

Results of Experiment 2 showed that the chlorophyll *a* concentration did not decrease compared to the control after maintaining the cultures desiccated at 10% HR for one week after a pre-treatment in BG11 with 0.2 M NaCl (Figure 4). This pre-treatment did not prevent the chlorophyll *a* to decrease after the rehydration + UVR treatment, however the concentration was significantly higher (0.4 µg · mL^−1^; Figure 3) than in absence of NaCl pre-treatment (0.03 µg · mL^−1^; Figure 3). The different pre-treatments did not seem to influence the carotenoids concentrations after desiccation, but they were again significantly higher in rehydrated-UVR exposed cultures pretreated with 0.2 M NaCl (Figure 3). Interestingly, in this experiment, we detected the production of scytonemin for the first time (Figure 3) after rehydrating and exposing the cultures to UVR (Figure 3). A significantly higher concentration was found in cultures preincubated with 0.2 M NaCl (0.31 µg · mL^−1^; Figure 3) compared to 0 M NaCl.

**Figure 4:**
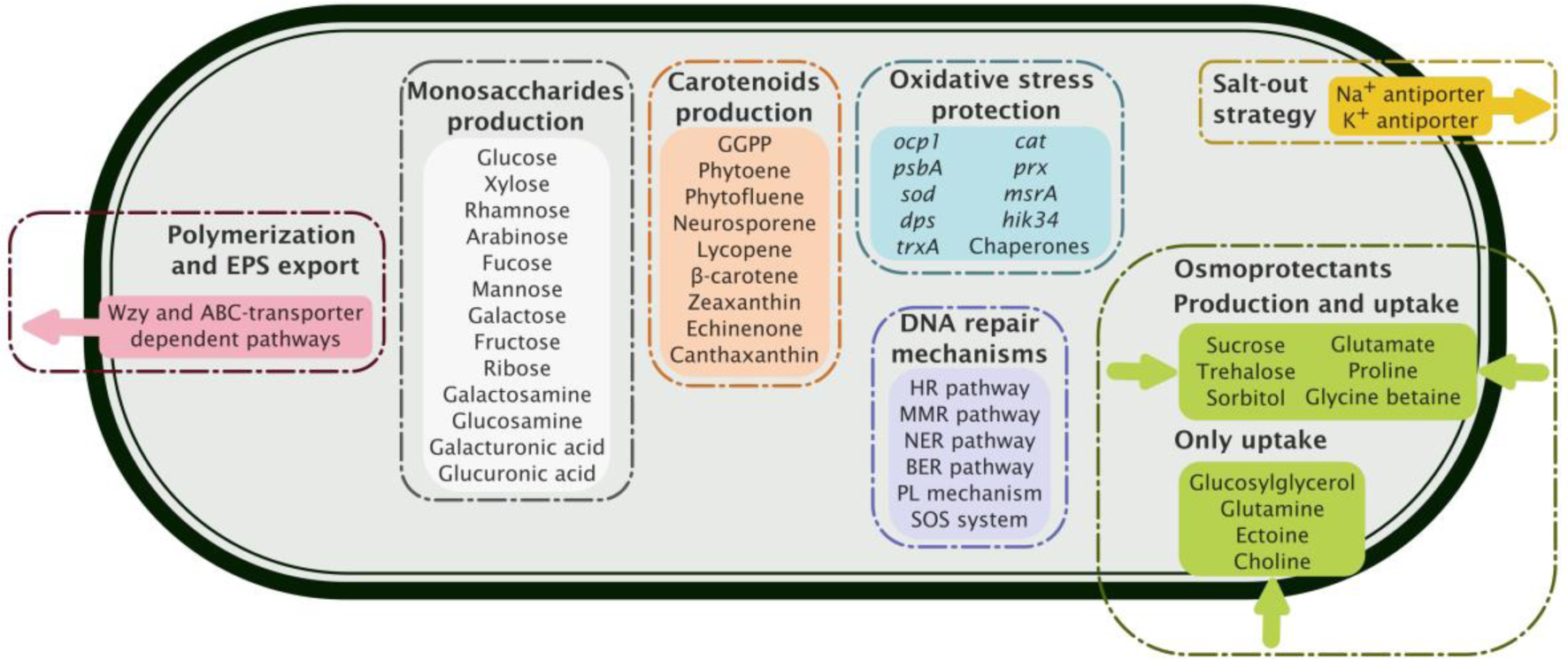
Complete pathways related to stress resistance mechanisms observed in ULC029

The production of exopolysaccharides increased after desiccation in the absence of a NaCl pre-treatment. However, it increased further after rehydration and UVR exposure whatever the pre-treatment (Figure 3). The total content of exopolysaccharides was always significantly lower when *S. frigidus* was pre-grown in medium with 0.2 M NaCl, except for the control (Figure 3). The loosely-bound fraction of EPS was significantly different between salinity pre-treatments (Suppl. Figure 5). This fraction was always significantly higher in cultures pre-treated with 0 M NaCl, even in the control treatment (Suppl. Figure 5). The only significant increase of EPS after desiccation compared to the control was found in the LB-EPS fraction of the 0 M NaCl pre-treatment (Suppl. Figure 5). After the rehydration + UVR treatment, both EPS fractions significantly increased compared to the control and desiccation treatments (Suppl. Figure 5).

Finally, the production of the osmoprotectants trehalose and sucrose was significantly higher after the desiccation and rehydration + UVR treatments in the absence of a NaCl pre-treatment, except for the control treatment where there were no differences (Figure 3). Their concentration significantly increased after desiccation compared to the control for both pre-incubation types, followed by a significant decrease after rehydration + UVR whatever the pre-treatment type (Figure 3). The highest concentrations were observed after desiccation of the strain pre-grown without NaCl, namely for sucrose (1130.3 mg ·mL^−1^), followed by trehalose (981 mg ·mL^−1^; Figure 3).

The efficiency of the photosynthesis (*F_v_/F_m_*) did not show differences when *S. frigidus* had been pre-cultured with or without 0.2 M NaCl in any treatment (Figure 3). However, the strain was photosynthetically inactive after desiccation (∼ 0; Figure 3), as expected, and started recovering the initial values of *F_v_/F_m_* after rehydration and UVR treatment (Figure 3).

### Mechanisms to cope with stress potentially observed in the genome of ULC029

#### Genome assembly

After sequencing and combining short and long reads of a culture of *S. frigidus* ULC029, we obtained 8 bins (Suppl. Table 4). One of them was identified by CheckM as a cyanobacterium, with a completeness of 99.29 % and a contamination of 0.35% (Table 1). This genome was assigned to *S. frigidus* ULC029 and has a total length of 6403461 bp and a GC% of 50.68 (Table 1). It is composed by 1 long contig (6320863 bp) and 2 short contigs (Suppl. Figure 6), with a coding ratio of 80.6 % (Table 1). Gene annotation by DFAST resulted in 5587 CDSs, 6 rRNAs and 47 tRNAs (Table 1). GTDBtk classification placed the genome in the *Stenomitos* cluster (see Suppl. Figure 6) with *Stenomitos frigidus* ULC018 (GCF_003003795.1), which shows the closest ANI value (79.22%) and was isolated from a neighboring lake microbial mat in the same Antarctic region (*Stenomitos* sp. ANT.LH53B.2; Taton et al. 2006).

**Table 1:** Genome characteristics obtained from results produced by Quast, DFAST, anvi’o and CheckM software.

|  | *Stenomitos frigidus*<br>ULC029 |
| --- | --- |
| Contigs | 3 |
| Largest contig (bp) | 6320863 |
| Genome size (bp) | 6403461 |
| GC (%) | 50.68 |
| N° of CDSs | 5,587 |
| N° of rRNA operons | 3 |
| N° of tRNA | 47 |
| N° of CRISPRS | 15 |
| Coding Ratio (%) | 80.6 |
| COGs assignation | 4042 |
| KEGGs assignation | 2026 |
| CheckM completeness (%) | 99.29 |
| CheckM contamination (%) | 0.35 |

#### Mechanisms to cope with osmotic, desiccation, and UVR stress

Genome annotation was enhanced after using the COG and KEGG databases, which resulted in 4042 and 2026 annotated genes, respectively (Suppl. Table 1). Among them, we identified 140 genes coding for proteins known to be involved in resistance mechanisms to desiccation, UVR and salinity stress (Suppl. Table 5): production of the EPS matrix, photoprotection, oxidative stress protection, DNA repair and osmotic stress protection. 113 of them were annotated with the COGs and/or KEGGs databases and 27 were identified using tblastn against the protein sequence of interest downloaded from NCBI (Suppl. Table 3).

Concerning the production of the EPS matrix, we detected genes coding for proteins related to the production of the following monosaccharides that could compose the EPS matrix: glucose (*glgAC*), xylose (*uxs*), rhamnose (*rfbABCD*), arabinose (*uxe*), fucose (*fcl*), mannose (*manA*), galactose (*galE*), fructose (*pgi*), ribose (*rpiA*), glucosamine (*glmU*), glucuronic acid (*ugd),* galactosamine (*wbpP*) and galacturonic acid (*cap1J;* Suppl. Tables 3 and 5). Though three pathways were described in cyanobacteria to assemble and export polysaccharides (Pereira et al. 2019), only two of them are complete in this genome: the Wzy (*wza*, *wzb*, *wzc*, *wzx* and *wzy*) and the ABC-transporter (*kpsTMED*) dependent pathways (Suppl. Tables 3 and 5). Two of the main proteins of the synthase dependent pathway could not be annotated: AlgE and AlgK, while Alg8 and Alg44 were found (Suppl. Table 5).

Concerning the photoprotection strategies of cyanobacteria, we focused on the production of pigments (carotenoids and scytonemin) and of mycosporine-like amino acids. The genes encoding enzymes involved in the production of most carotenoids were annotated: *crtBEHOPQRW* and *cruA* (Suppl. Table 5). Therefore, the list of carotenoids that *S. frigidus* ULC029 could potentially produce include at least: geranylgeranylpyrophosphate (GGPP), phytoene, phytofluene, neurosporene, lycopene, β-carotene, zeaxanthin, echinenone and canthaxanthin (Suppl. Table 5). Concerning the synthesis pathway of scytonemin, we could not find all the proteins and genes involved. The annotated genes composing the scytonemin operon (Klicki et al. 2018) were: *trpABCDEFG* and *tyrA*, which are involved in the biosynthesis of tryptophan and p-hydroxyphenylpyruvic acid, respectively; and *aroBG*, involved in the shikimic acid pathway (Suppl. Table 5). The genes coding for proteins involved in the last steps of the scytonemin biosynthesis, *scyABDE*, were assigned by tblastn (Suppl. Tables 3 and 5), except for the gene *scyC*. Besides, the genes belonging to the *ebo* cluster, involved in the translocation of scytonemin to the periplasm, were not annotated (Suppl. Table 3). Finally, all the genes (*mysABE)* coding for enzymes involved in the synthesis of MAAs were assigned by tblastn, except for *mysC*, which encodes the protein producing mycosporine-glycine, the branching point for the rest of MAAs.

Several oxidative stress protection mechanisms were identified in the ULC029 genome. First, the gene *ocp1*, coding for the orange-carotenoid-binding protein, involved in nonphotochemical quenching and thermal dissipation of high light energy (Chrismas et al. 2018), was assigned by tblastn (Suppl. Tables 3 and 5). Another annotated gene was *psbA* (Suppl. Table 5), which codes for a chlorophyll-binding protein (D1) of PSII. It is induced by light stress and would confer oxidative stress protection to the photosynthetic system (Kopf et al. 2015). One of the most known mechanisms against reactive oxygen species, the superoxide dismutase (*sod*), was also annotated (Suppl. Table 5), specifically, the combination of Fe and Mn SOD, which is the type found in filamentous nonheterocystous cyanobacteria (Napoli et al. 2021). Other genes related to oxidative damage protection that were detected are: a methionine sulfoxide reductase (*msrA*), the Mn containing catalase *(cat*), the DNA-binding protein (*dps*); and peroxiredoxin (*prx*). Besides, several genes coding for molecular chaperones involved in protein folding were annotated too: *dnaKJ*, *groEl/ES*, *htpG* and *clpB*, as well as two of their regulatory factors, *hik34* and *trxA* (Suppl Table 5).

We identified four DNA repair mechanisms in the genome, as shown in Suppl. Table 5. First, the homologous recombination pathway, composed by the genes: *holB* (DNA polymerase III), *recQG* (ATP-dependent DNA helicase), *recJ* (single-stranded-DNA-specific exonuclease), *ssb* (single-strand DNA-binding protein), *recORF* (recombinational DNA repair proteins), *recA* (recombinase), *uvrABC* (hollyday junction enzymes) and *priA* (primosomal protein).The genes *holB*, *recJ* and *ssb*, are also involved in the mismatch repair (MMR) pathway, together with the genes *lig* (DNA ligase), *uvrD* (DNA helicase), *mutSL* (ATPases), *exoVII* (exonuclease VII) and *dam* (site-specific DNA-adenine methylase). The nucleotide excision repair (NER) pathway was also fully detected: the already mentioned *uvrABCD* and *lig*, *dpoI* (DNA polymerase I) and *mfd* (transcription-repair coupling factor). The base excision repair (BER) pathway is also complete in the genome, being composed by: *fpg* (nucleotide glycosylase), *nei* (endonuclease VIII), *nth* (endonuclease III), *xth* (exonuclease III), *mpg* and *alkA* (DNA-3-methyladenine glycosylase and II), *mutY* (A/G-specific adenine glycosylase), *udg* (uracil-DNA glycosylase), as well as *lig* and *dpoI*. Other genes found that are involved in DNA repair mechanisms were *pL*, coding for a DNA photolyase which corrects DNA damage caused by solar light, and *lexA*, a transcriptional repressor involved in the regulation of the SOS repair system.

Concerning the mechanisms for osmotic stress protection, we focused on the salt-out strategy and the osmoprotectants production and uptake (Suppl. Table 5). The genes coding for the Na^+^ and K^+^ antiporters (*nhA* and *ktrAB*, respectively) and the *ktrE* gene (that contributes to the import of K^+^ into the cells), which are involved in the salt-out strategy, were detected in the genome. However, only one member of the *kdp* gene family (*kdpD*), related to the import of H^+^ into cells, was detected. *kdpD* is involved in the activation of the rest of *kdp* genes. The second resistance strategy in highly saline environments was the synthesis and/or uptake into the cells of osmoprotectants. *S. frigidus* ULC029 has the main genes involved in the production of the following ones: sucrose (*spp* and *sps*), trehalose (*treYZX* and *glgABC*), sorbitol (*srlD*), glutamate (*gltS*), proline (*proABC*), glycine betaine by the choline pathway (*betAB*) and the glycine pathway (*GNMT* and *DMT*). However, this strain would not be capable of producing glycosylglycerol, as both genes coding for the enzymes involved in its production pathway are missing (*ggpS* and *ggpP* TS4), nor ectoine because the gene coding for the L-ectoine synthase (*ectC*) was not found (Suppl Table 3). The mechanisms for osmoprotectant uptake that were detected are: ABC-type sucrose/trehalose/glucosylglycerol transport system (*ggtA*), ABC transporter for glutamine and glutamate (*glnHPQ* and *gluA*), neutral aa transporter N-I type (*natABCDE*) and one of the genes required for the N-II type transporter of acidic and polar aa (*natF*), glycine betaine and proline transporter (*opuABC*), and ABC transporter for ectoine, glycine betaine, proline and choline uptake (*proXWV*).

## Discussion

The strain ULC029 belongs to a cluster of 16S rRNA sequences with two lineages, one containing Antarctic strains (Taton et al 2006; Rego et al. 2019) and one with strains from Spanish arid deserts and a dry Siberian steppe (Temraleeva, 2018; Roncero-Ramos et al. 2019) (Suppl. Figure 2). The similarity between the sequences of ULC029 and CAU10 is 99% (1367 positions), which may indicate that they belong to the same species and underlines the relevance of this experimental data for soil restoration strategies. Although taxonomic uncertainties and the recent establishment of the genus *Stenomitos* complicate the comparison with literature data, it seems probable that similar resistance mechanisms are present in the thin filamentous genera from the Leptolyngbyaceae (Strunecký et al. 2022).

Several resistance mechanisms to abiotic stress factors have been already reported for strains belonging to the genus *Leptolyngbya*, in which ULC029 was previously included, such as: the production of photoprotective pigments, carotenoids and scytonemin (Lin and Wu 2014; Kokabi et al. 2019); the production of mycosporine-like amino acids and the identification of the related genes (Rossi et al. 2012; Shimura et al. 2015); the production of trehalose (Shimura et al. 2015), proline (Lin and Wu 2014), sucrose, glycogen and glucosylglycerol (Keshari et al. 2019) as compatible solutes during events of osmotic stress or desiccation; and the increase of the activity of the ROS-scavenger enzymes peroxidase and superoxide dismutase after drought stress (Lin and Wu 2014). In this study, we analyzed the ecophysiological response to osmotic, desiccation and UVR stress and the potential response of *S. frigidus* ULC029 by stress-genes mining. Moreover, we provide a high-quality genome for this strain, being the second one identified as *S. frigidus* that is available in public databases.

### The response of *S. frigidus* to osmotic stress and UVR exposure

We observed that *S. frigidus* ULC029 cannot tolerate NaCl concentrations higher than 0.7 M and its optimal NaCl concentration is between 0.05 to 0.2 M (Figure 2). Thus, it belongs to the cyanobacterial group with low tolerance to salinity (Mackay et al. 1984). We also show that after one month of cultivation in BG11 medium with 0.2 and 0.7 M NaCl, *S. frigidus* ULC029 shows signs of a high stress (Suppl. Figure 4). However, it is resilient by showing a full recovery of the *F_v_/F_m_* values after the osmotic stress ceased (recovery phase, Suppl. Figure 4). Moreover, although a decrease in chlorophyll *a* concentration is generally observed at all salinities and after three days of exposure to UVR, cultures growing in media with 0.2 M NaCl show the highest chlorophyll *a* concentration after UVR (Figure 2c). However, they also had a higher stress than cultures growing in 0 and 0.05 M NaCl (Suppl. Figure 4). Carotenoids concentrations show a similar pattern to chlorophyll *a* concentrations (Figure 2), which is related to the production of primary carotenoids, those bound to the photosystems and participating in the photosynthesis process (Mulders et al. 2015; Rehakova et al. 2019). According to its genome, the main primary carotenoids that *S. frigidus* ULC029 is able to produce are the zeaxanthin and echinenone (Suppl Table 5; Figure 4). However, it has been shown (Mulders et al. 2015; Rehakova et al. 2019) that primary carotenoids are usually degraded under stress and replaced by secondary carotenoids that are involved in photoprotection, such as canthaxanthin or β-carotene, for which pathways are complete in ULC029 genome as well (Suppl. Table 5; Figure 4). Therefore, the carotenoid composition before and after UV radiation could be different with increasing concentrations of secondary carotenoids after UVR. Finally, as the addition of 0.2 M NaCl was stressing the strain but it could still survive and grow, we selected this concentration to further explore the influence of osmotic stress on the cyanobacterial resistance to desiccation and UVR in a second experiment.

### Performance of *S. frigidus* under multiple stress factors

In Experiment 2, after 4 months pre-adapting ULC029 to the osmotic stress induced by 0.2 M NaCl, there were no significant differences in the variables measured compared to non pre-treated cultures (Control treatment, Figure 3). Moreover, this hardening pre-treatment by osmotic stress did activate resistance mechanisms that allowed ULC029 to withstand better the subsequent desiccation and UVR treatments compared to the non pre-adapted ones, as it is shown by a significant higher concentration of chlorophyll *a* and similar *F_v_/F_m_* values (Figure 3).

The EPS production increased significantly after desiccation, rehydration and UVR (Figure 3), which is in accordance with previous works (Wu et al. 2021; Shang et al. 2019; Li et al. 2022). Given the genome annotation results (Suppl. Tables 3 and 5), the two pathways that are complete and could be triggered to assemble and export EPS are the Wzy and the ABC-transporter dependent pathways (Figure 4). Besides, according to the genome annotation, the EPS matrix could be composed by glucose, xylose, rhamnose, arabinose, fucose, mannose, galactose, fructose, ribose, glucosamine, glucuronic acid, galactosamine and galacturonic acid. The possibility of producing an EPS matrix with such a complex composition of monosaccharides could also favor the colonization of the inoculated soils by a more diverse microbial community (Chamizo et al. 2019). Also, most of these sugars have been found in artificially induced biocrusts (Román et al. 2021) and some of them have been related with a protective role under desiccation, such as the galacturonic acids (Tamaru et al. 2005). In spite of the increase in EPS production after desiccation and UVR stress, the hardening treatment by osmotic stress coincided with a significantly lower EPS concentration compared to non-pre-adapted cultures (Figure 3). This is in line with the controversy already reported by others (Cruz et al. 2020) about the increasing and decreasing evolution of EPS production found in different cyanobacteria cultivated with NaCl.

Concerning the production of osmoprotectants, this strain was capable of significantly increasing the concentration of both sucrose and trehalose after one week under high dehydration stress (Figure 3), as previously shown by others (Raanan et al. 2016). Sucrose concentration values were slightly higher than the trehalose ones in most cases (Figure 3), suggesting that sucrose might have a more important role in the response to stress than trehalose in this cyanobacterium. Both osmoprotectants would be acting against cellular osmotic unbalance due to dehydration (Potts, 1994), by preventing protein denaturation (Borges et al. 2002) and stabilizing membrane structure (Hincha and Hagemann, 2004). After three days of rehydration and UVR exposure (Figure 3), the concentration of both osmoprotectants significantly decreased. Fagliarone et al. (2020) showed that *Chroococcidiopsis* sp. desiccated cells under UVR could maintain a basal number of mRNA genes corresponding to the sucrose and trehalose production pathways. This suggests that if we had exposed *S. frigidus* to UVR in a desiccated instead of a rehydrated state, the concentration of sucrose and trehalose might have been higher. Moreover, the decrease in osmoprotectants was stronger in pre-treated cultures (0.2 M NaCl) with concentrations values similar to those before desiccation (Control; Figure 3). Besides, contrary to previous works (Keshari et al. 2019), the accumulation of both osmoprotectants in all pre-treated cultures was lower than in those not pre-treated (0 M NaCl; Figure 3). This has been previously shown in *Anabaena fertilissima* (Swapnil and Rai, 2018), where the production of sucrose and trehalose increased with NaCl concentrations, until a maximum value was reached (276 and 280%, respectively, of control) and the production started to decrease. These authors also tested the addition of other molecules to cause osmotic stress, such as Na_2_SO_4_, which showed a lower toxicity than NaCl and induced a higher production of sucrose and trehalose. Therefore, using different molecules to induce osmotic stress in the pre-adaptation medium could help increasing the production of these osmoprotectants in *S. frigidus* as well. Concerning the genome annotation, *S. frigidus* ULC029 shows all the genes required for the production and/or uptake of trehalose and sucrose but also of sorbitol, glucosylglycerol, glutamate, proline, ectoine and glycine betaine (Suppl. Table 5; Figure 4). The lower production of sucrose and trehalose under osmotic stress (Figure 3) might be counteracted in *S. frigidus* ULC029 by the synthesis of any of these other osmoprotectants. However, the ability to accumulate glucosylglycerol and glycine betaine is normally considered as an indicator of an haloterant or halophilic cyanobacterium (Pade and Hagemann, 2014), which is not the case of ULC029, as it was not able to tolerate NaCl concentrations higher than 0.7 M (Figure 2). This deviation from the classical pattern has already been shown in marine cyanobacterial species lacking the genes for glucosylglycerol, such as *Crocosphaera watsonii* and *Trichodesmium erythraeum* (Pade and Hagemann, 2014). This supports the importance of enlarging the cyanobacterial genome databases and suggests the need for revising the cyanobacterial classification by osmotic resistance based on the type of osmoprotectant produced, starting by performing metabolomics analyses to determine if these genes are expressed after the exposition to different stress factors, such as salinity.

Finally, both pigments measured (carotenoids and scytonemin concentrations) showed significantly higher values after desiccation and UVR exposition in cultures pre-treated with NaCl (Figure 3). Our hypothesis is that in Experiment 1, the carotenoids probably mostly consist of secondary carotenoids synthetized after desiccation and UVR stress treatments. In spite of the significant higher concentration of carotenoids in cultures pre-treated with NaCl in Experiment 2, the carotenoids concentration decreased in both cases after desiccation and UVR treatments (Figure 3). These results are in contrast to previous works which showed an increase of carotenoids concentration after UV radiation (Kokabi et al. 2019) and desiccation (Li and Wu 2014) in strains belonging to the *Leptolyngbya* genus. Carotenoids would serve as photoprotectants and quenchers (Llewellyn et al. 2020). The decrease of primary carotenoids might be related to the decrease in chlorophyll *a*, as they are involved in the photosynthesis process and are usually degraded under stress (Mulder et al. 2015; Rehakova et al. 2019). Li and Wu (2014) discussed that only drought-tolerant cyanobacteria showed an increase in carotenoids after desiccation. As *S. frigidus* ULC029 is a freshwater cyanobacteria, its production of secondary carotenoids might not be activated under desiccation stress, which could explain the unexpected decrease of total carotenoids after desiccation and UVR stress.

Interestingly, the production of scytonemin was detected by the extraction method used after the desiccation and UVR treatments in both cultures in Experiment 2 (Figure 4). In contrast, there was no production of scytonemin in Experiment 1 after the exposition to UVR. However, after the desiccation and UVR treatments in Experiment 2, it increased in both cultures and was significantly higher in the strain pre-adapted with 0.2 M NaCl (Figure 4). Thus, it seems that a previous desiccation step could be necessary to trigger the production of scytonemin in this strain, and it could be also enhanced by adding a preliminary osmotic stress. Desiccation and osmotic stress in fact trigger a similar physiological response in cyanobacteria (Billi and Potts 2002), especially during the early stages of air drying. It has been previously shown (Fleming and Castenholz 2007; Dillon et al. 2002) that the scytonemin production is affected by periodic expositions to desiccation, which is in agreement with our results, and could be explained by its probable role in the stabilization of the EPS matrix (Gao 2017). On the other hand, to our knowledge, it is the first time that an increase in scytonemin production linked to a preliminary osmotic stress has been found. Previous works could not detect this relation, i.e. Bennett and Soule 2022 showed no activation in *Nostoc punctiforme* ATCC 29133 of the genes related to scytonemin production after osmotic stress, similarly to our results in Experiment 1. Thus, this supports the hypothesis that scytonemin production can be induced by a more generalized environmental stress in cyanobacteria (Rehakova et al. 2019; Dillon et al. 2002). However, after analyzing the genome annotation, we could not find all genes related to scytonemin production, specially *scyC* and the *ebo* cluster (Suppl. Table 3). The former gene is involved in a key step of scytonemin biosynthesis (Ferreira et al. 2016) and the latter in the translocation of scytonemin to the periplasm (Klicki et al. 2018). This might be based on gaps due to the genome assembly process or the removal of these genes during the genome assembly, because there are usually several repeats (Zhou et al. 2014).

### Future prospects for cyanobacterial hardening

Previous works (Giraldo-Silva et al. 2019a, 2020; Román et al. 2021b) already showed that a hardening pre-treatment could improve soil cyanobacteria growth in field conditions, using chlorophyll *a* concentration as a proxy. Our work corroborates these previous results for *S. frigidus* ULC029 but using a multiple stress hardening treatment (Figure 4). The cultivation of ULC029 with a NaCl concentration inducing osmotic stress, but allowing its growth (Figure 2), induced a higher resistance to desiccation and UVR. This was shown by similar or higher values of chlorophyll *a* concentration in pre-treated cultures compared to cultures without osmotic stress (Figure 4). The resistance mechanisms involved were not identified by our ecophysiological measurements, as neither EPS, sucrose or trehalose concentrations were higher in cultures under osmotic stress compared to controls (Control; Figure 4). However, the genome annotation shows the capacity of this strain to produce many other osmoprotectants, enzymes related with oxidative stress (such as the superoxide dismutase (SOD), the orange carotenoid protein (OCP) or the chlorophyll-binding protein (D1) or several DNA repair pathways (Suppl. Tables 3 and 5; Figure 5) which were not analysed during our experiments and could have been activated by osmotic stress. On the other hand, the exposure of *S. frigidus* ULC029 to UVR after desiccation (Figure 4) or without this step (Figure 2), shows that the production of scytonemin is triggered, especially under osmotic stress (Figure 4). Moreover, EPS production was significantly enhanced after desiccation and UVR exposure in both cases (Figure 4). In view of these results, we propose a hardening treatment based on the application of sequential stress factors to the cultures, which should always include at least one stress factor related to osmotic unbalance (osmotic or desiccation stress), and if UVR is applied, it should be more effective afterwards. However, more studies analysing the response of cyanobacteria to multiple stress factors are required to fully understand their mechanisms to survive in extreme environments, and thus, to be applied in nature-based biotechnological tools for soil restoration or as biofertilizers in agriculture. In addition, the number of cyanobacterial genomes in public databases with high completeness and low contamination values is low (Cornet et al. 2018), being only seven to date if we only consider polar strains (Pessi et al. 2023). Besides, none of these genomes belong to the genus *Stenomitos*, which is closely related to *Leptolyngbya* sp. BC1307 (Chrismas et al. 2018). The release of more high-quality genomes would improve our knowledge of the resistance mechanisms that could be potentially activated under stress, as well as enable the application of transcriptomic analyses to monitor the cyanobacterial response to stress.

## Conclusions

More efficient and effective hardening methods of cyanobacterial inocula are required to improve biocrust restoration in field. In this work, we tested a feasible and easy-to-do hardening method for *Stenomitos frigidus* ULC029 based on a pre-treatment by adding salt to the culture medium to induce osmotic stress. This method allowed the strain to maintain its chlorophyll *a* concentration under a severe desiccation event, followed by rehydration with UVR exposition. The latter induced an expected decrease of chlorophyll *a* concentration values which were, however, slightly higher in cultures under osmotic stress. Here, we show that the sequential application of multiple stress factors to cyanobacterial cultures induces the production of three of the main mechanisms to survive in dryland soils: scytonemin, carotenoids and EPS. Besides, we provide a new high-quality genome for this Antarctic strain, which could be used in future studies to deepen our knowledge on its resistance mechanisms by, i.e., performing transcriptomics.

## Author contributions

B.R.R., A. W. and V.S. conceived the experiments. B.R.R. performed most of the analyses. B.D. gave support with bioinformatic analyses. A.R. and I. V. contributed analysing the osmoprotectant contents. B.R.R. wrote the manuscript. B.R.R. and A. W. found the financial support. All authors provided important feedbacks and contributed to the writing of the manuscript.

## Acknowledgments

BRR was supported by the IPD-STEMA Programme and the Special Funds for Research (R.DIVE.0899-J-F-I, University of Liège), and by the Junta de Andalucía (PAIDI-DOCTOR 21_00571), VS and BD by the PhD FRIA fellowship from the FRS-FNRS, and AW is Senior Research Associate of the FRS-FNRS. We would like to thank Anne Catherine Ahn and Kim Beets for their help with the cultivation of the strain of the BCCM/ULC cyanobacterial culture collection, Emmanuel Mignolet for the building of the UV box, Elie Verleyen for borrowing the UV sensor, and Luc Cornet for discussion of the genomic data.

## Supplementary Material

**Suppl. Table 1:** KEGG and COG genes annotation results can be found in Supplementary Material II file.

**Suppl. Table 2:** Gene sequences of ULC029 genome can be found in Supplementary Material II file.

**Suppl. Table 3:** The queries used to manually search genes using tblastn against ULC029 genome (e-value threshold of 1e-5 and a score > 50 bits) can be found in Supplementary Material II file.

**Suppl. Table 4:** CheckM and QUAST results of all bins obtained.

**Suppl. Table 5:** Proteins found in the genome classified by mechanisms associated with resistance to different stress factors. Each row includes the function and gene of each protein, as well as the annotation results from the three methods used (COG and KEGG database, or the tblastN query alignment).

**Suppl. Figure 1:** A picture of *Stenomitos frigidus* ULC029

**Suppl. Figure 2:** Phylogenetic tree based on 16S rRNA gene sequences (1388 bp) obtained by the Maximum likelihood method, using the K2+G+I model (Kimura, 1980), calculated with MEGA X (Kumar et al. 2018). The bootstrap percentage of trees in which the associated taxa clustered together is shown next to the branches. The tree is drawn to scale (0.020 bar), with branch lengths measured in the number of substitutions per site. *S. frigidus* ULC029 sequence is highlighted by a red square and a red circle indicates the ancestral node of the cluster containing ULC029 and 6 strains, which origin is depicted by different symbols.

**Suppl. Figure 3:** Monitoring of temperature and relative humidity inside the desiccation chamber during Experiment 2.

**Suppl. Figure 4**: Photosynthetic efficiency (*F_v_/F_m_*) of *S. frigidus* ULC029 cultured in increasing concentrations of NaCl in function of incubation time until day 32. Afterwards, results of two parallel treatments are shown: 3 additional days of UVR exposure, and 33 additional days of recovery in BG11 medium.

**Suppl. Figure 5:** Comparison of the influence on Loosely and Tighly bound EPS content of desiccation and later rehydration steps while exposed to UVR on *S. frigidus* ULC029 cultured in BG11 with 0.2 M NaCl compared to the control. All data are means (n = 4). The error bars represent ±SE and letters indicate significant differences (p < 0.05) among treatments.

**Suppl. Figure 6:** Circular plot of *S. frigidus* ULC029 genome. Rings are as follows (outer-inner): annotated coding regions (CDs; yellow); open reading frames (ORFs; green); contigs (grey); GC content (black) and GC skew (orange and purple, for + and – skew, respectively). Doubled rings (yellow and grey) corresponds to both DNA strands.

## Notes

### Competing Interest Statement

The authors have declared no competing interest.

