## Supplementary material for "Ecophysiological and genomic approaches to cyanobacterial hardening for soil restoration": Supplementary Material.docx

**Running title: Cyanobacterial resistance to stress**

*Roncero-Ramos Beatriz^2^*

InBios-Molecular Diversity and Ecology of cyanobacteria, University of Liège, 4000 Liège, Belgium

Department of Plant Biology and Ecology, University of Seville, 41012 Seville, Spain

*Savaglia Valentina*

InBios-Molecular Diversity and Ecology of cyanobacteria, University of Liège, 4000 Liège, Belgium

Laboratory of Protistology & Aquatic Ecology, Ghent University, 9000 Ghent, Belgium

*Durieu Benoit*

InBios-Molecular Diversity and Ecology of cyanobacteria, University of Liège, 4000 Liège, Belgium

*Van de Vreken Isabelle*

TERRA-Biomass and Green Technologies, University of Liège, 5030 Gembloux, Belgium

*Richel Aurore*

TERRA-Biomass and Green Technologies, University of Liège, 5030 Gembloux, Belgium

*Wilmotte Annick*

InBios-Molecular Diversity and Ecology of cyanobacteria, University of Liège, 4000 Liège, Belgium

**Suppl. Figure 1:** A picture of *Stenomitos frigidus* ULC029 in optical microscopy

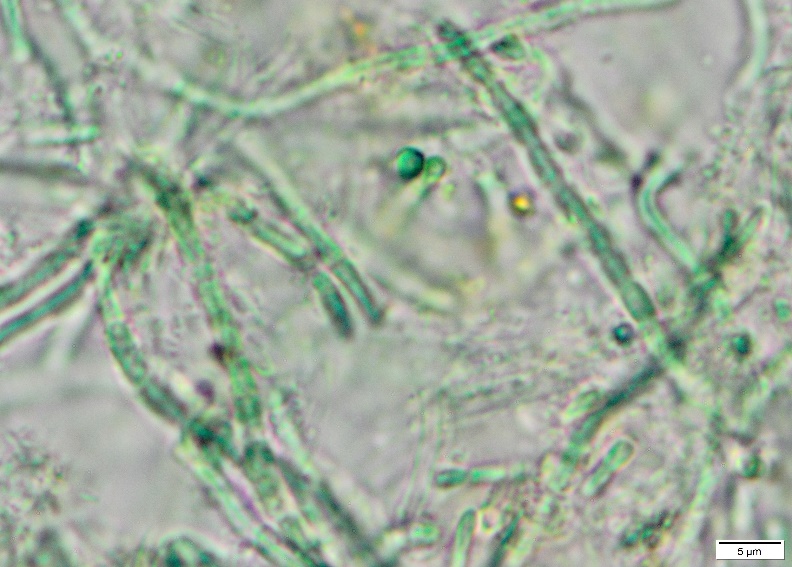

**Suppl. Figure 2:** Phylogenetic tree based on 16S rRNA gene sequences (1388 bp) obtained by the Maximum likelihood method, using the K2+G+I model (Kimura, 1980), calculated with MEGA X (Kumar et al. 2018). The bootstrap percentage of trees in which the associated taxa clustered together is shown next to the concerned nodes. The tree is drawn to scale (0.020 bar), with branch lengths measured in the number of substitutions per site. *S. frigidus* ULC029 sequence is highlighted by a red square and a red circle indicates the ancestral node of the cluster containing ULC029 and 6 strains, which origin is depicted by different symbols.

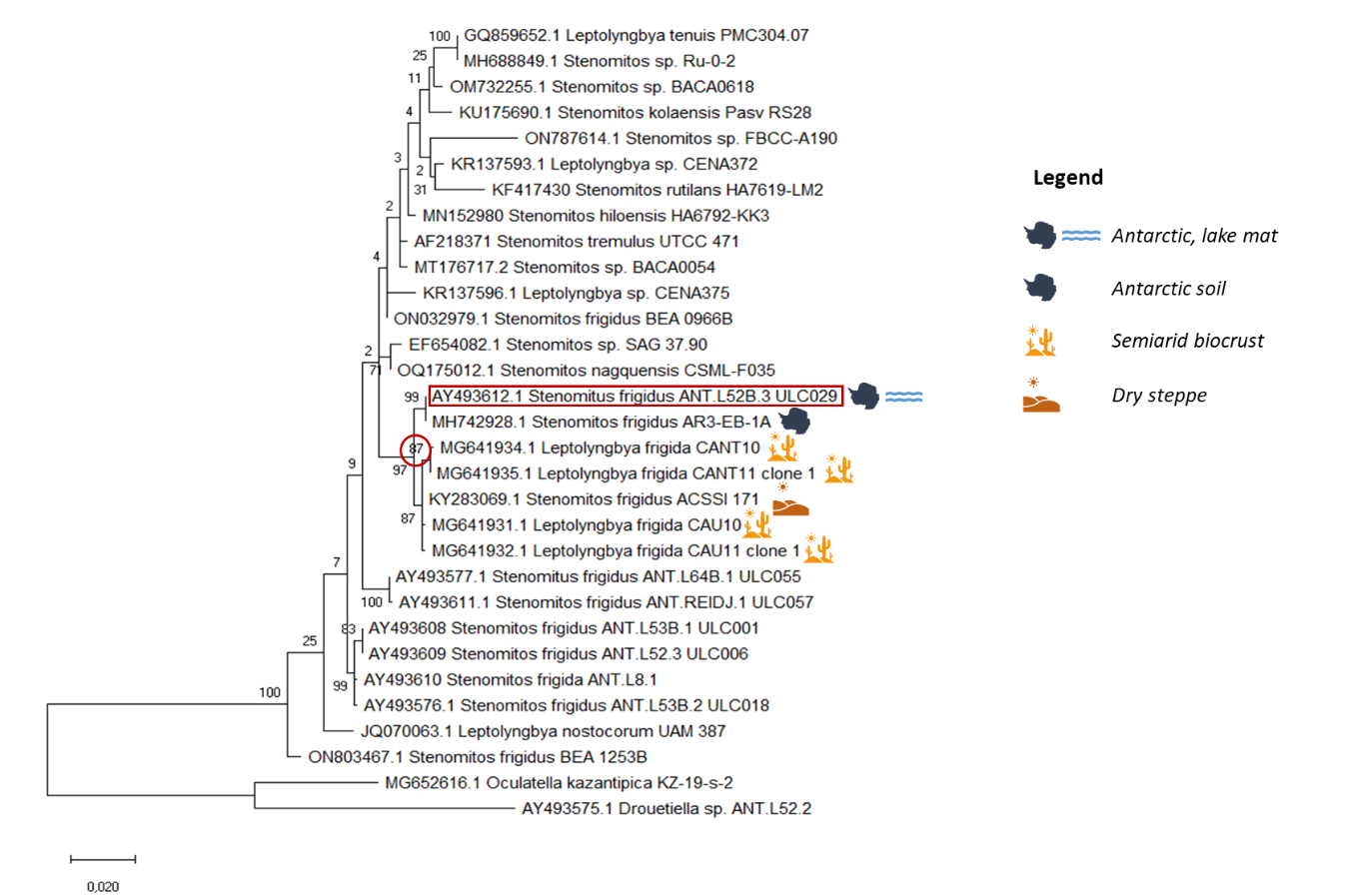

**Suppl. Figure 3:** Monitoring of temperature and relative humidity inside the desiccation chamber during Experiment 2.

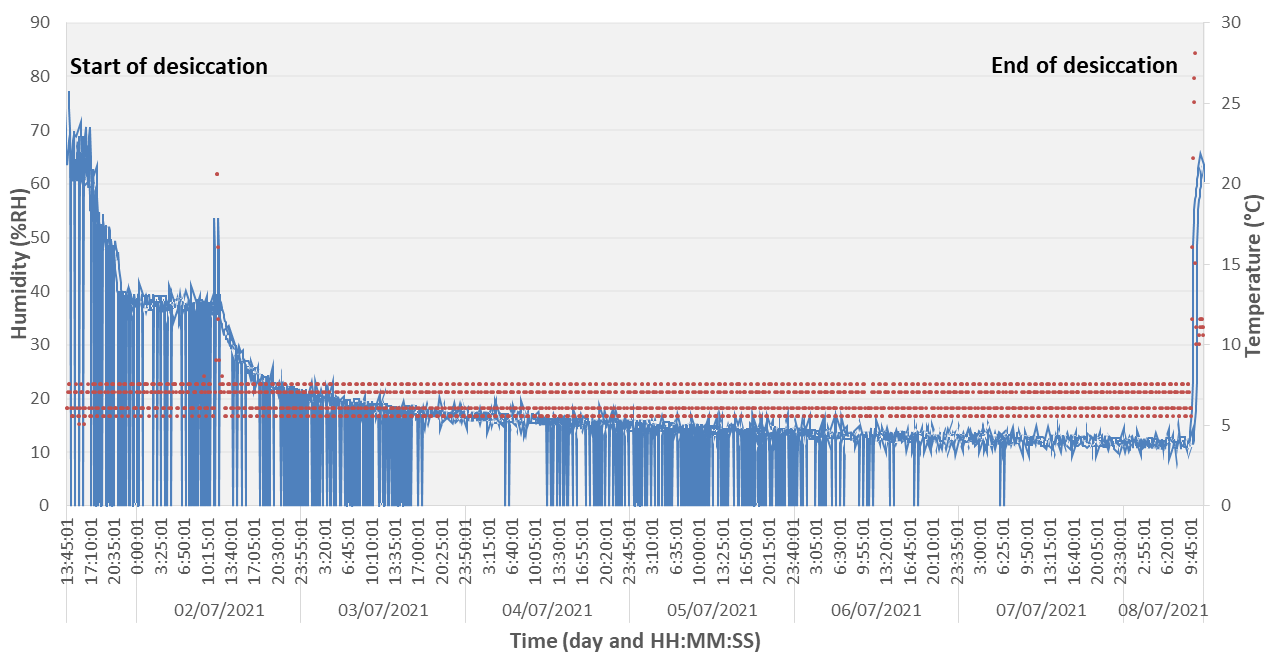

**Suppl. Figure 4**: Photosynthetic efficiency (*F_v_/F_m_*) of *S. frigidus* ULC029 cultured in increasing concentrations of NaCl in function of incubation time until day 32. Afterwards, results of two parallel treatments are shown: 3 additional days of UVR exposure, or 33 additional days of recovery in BG11 medium.

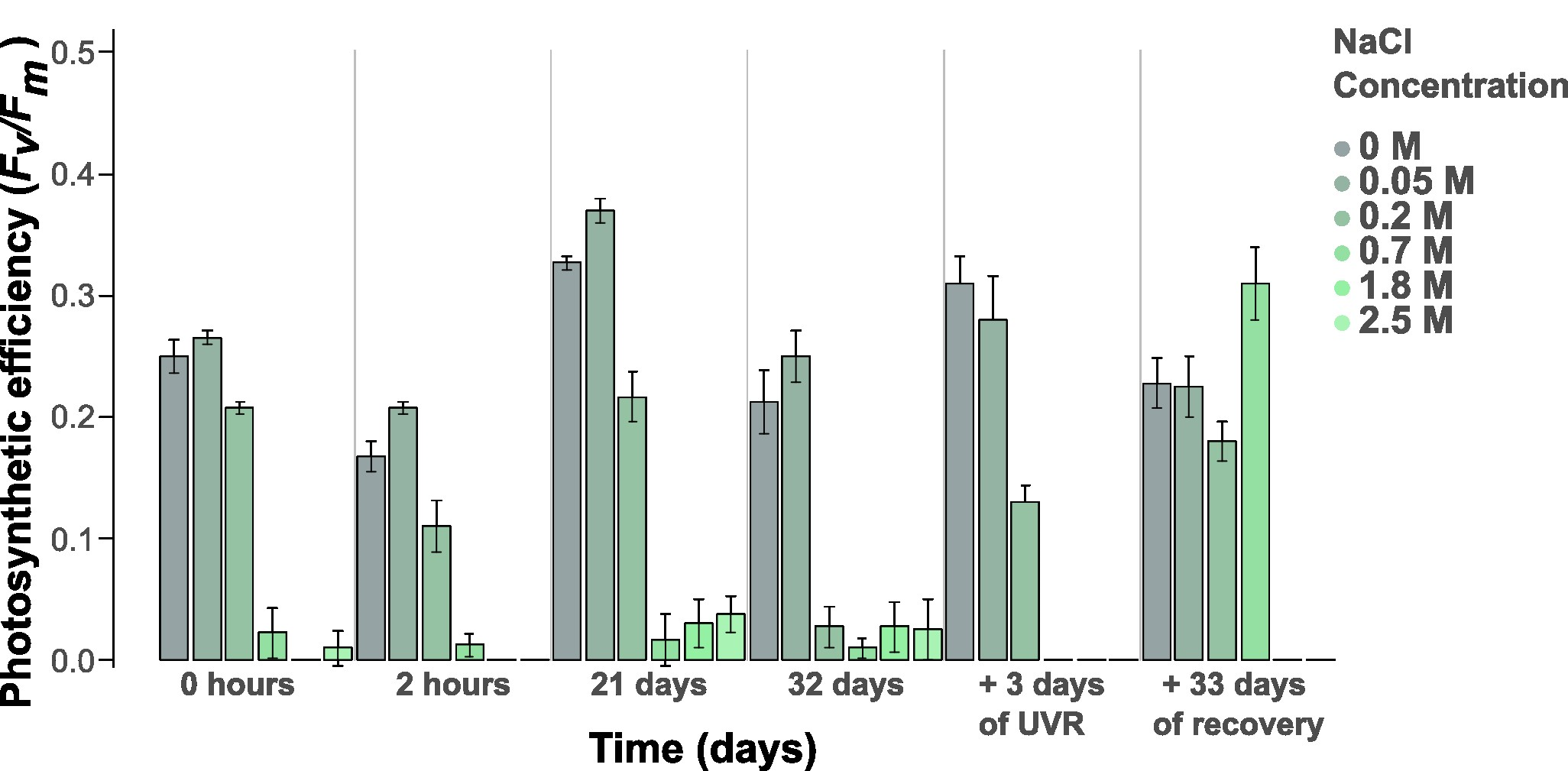

**Suppl. Figure 5:** Comparison of the influence on Loosely and Tighly bound EPS content of desiccation and later rehydration steps while exposed to UVR on *S. frigidus* ULC029 cultured in BG11 with 0.2 M NaCl compared to the control. All data are means (n = 4). The error bars represent ±SE and letters indicate significant differences (p < 0.05) among treatments.

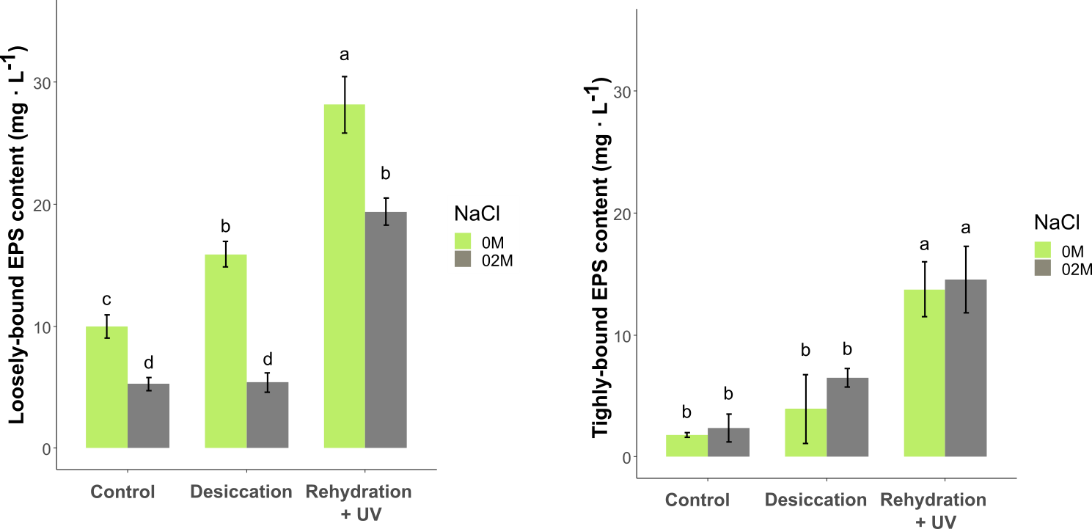

**Suppl. Figure 6:** Circular plot of *S. frigidus* ULC029 genome. Rings are as follows (outer-inner): annotated coding regions (CDs; yellow); open reading frames (ORFs; green); contigs (grey); GC content (black) and GC skew (orange and purple, for + and – skew, respectively). Doubled rings (yellow and grey) corresponds to both DNA strands.

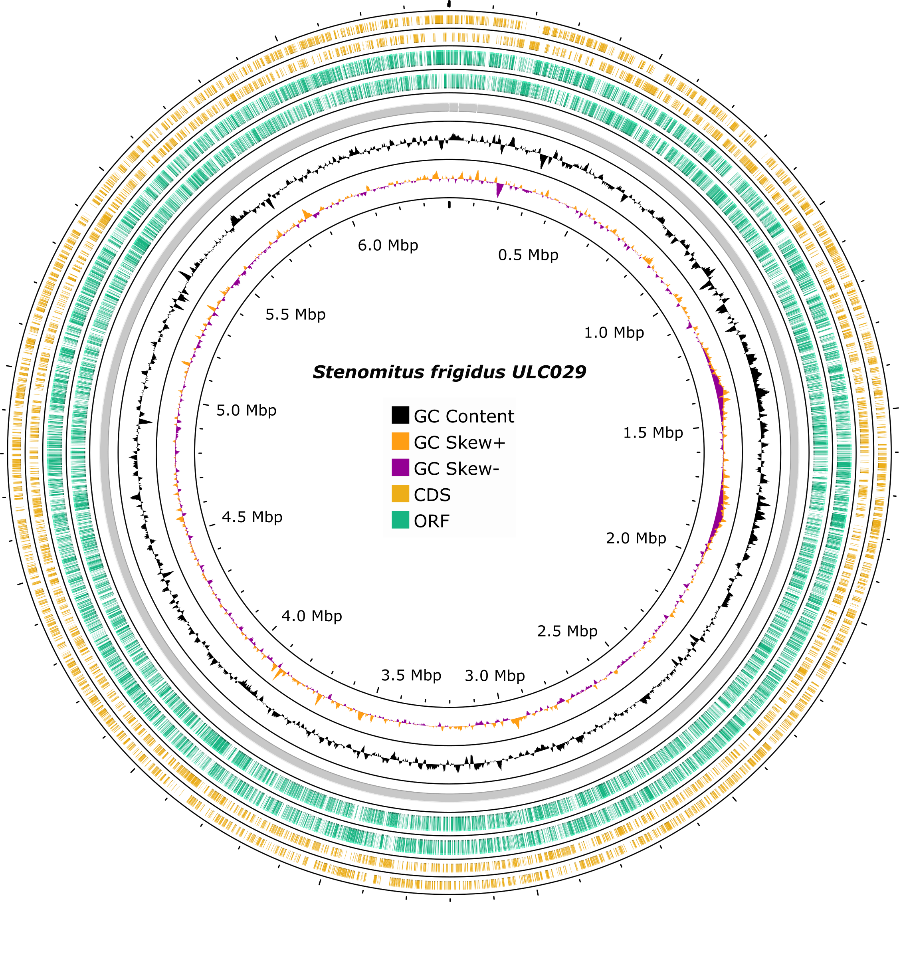

**Suppl. Table 1:** KEGG and COG genes annotation results can be found in Supplementary Material II file.

**Suppl. Table 2:** Gene sequences of ULC029 genome can be found in Supplementary Material II file.

**Suppl. Table 3**: The queries used to manually search genes using tblastn against ULC029 genome (e-value threshold of 1e-5 and a score > 50 bits) can be found in Supplementary Material II file.

**Suppl. Table 4**: CheckM and QUAST results of all bins obtained

| **Bin ID** | **Marker lineage** | **Size (bp)** | **Nº contigs** | **Completeness** | **Contamination** |
| --- | --- | --- | --- | --- | --- |
| **Bin 0** | root (UID1) | 360801 | 6 | 0 | 0 |
| **Bin 1** | p__Cyanobacteria (UID2182) | 6403461 | 3 | 99.29 | 0.35 |
| **Bin 2** | root (UID1) | 2648 | 1 | 0 | 0 |
| **Bin 3** | k__Bacteria (UID203) | 1242089 | 15 | 3.16 | 0 |
| **Bin 4** | f__Xanthomonadaceae (UID4214) | 3155256 | 1 | 96.93 | 0.45 |
| **Bin 5** | root (UID1) | 88804 | 3 | 0 | 0 |
| **Bin 6** | o__Burkholderiales (UID4000) | 3701889 | 36 | 49.36 | 0.03 |
| **Bin 7** | o__Rhizobiales (UID3449) | 3627737 | 43 | 13.43 | 0 |

**Suppl. Table 5**: Proteins found in the genome classified by mechanisms associated with resistance to different stress factors. Each row includes the function and gene of each protein, as well as the annotation results from the three methods used (COG and KEGG database, or the tblastN query alignment).

| Production of the EPS matrix | | | | | | | | |
| --- | --- | --- | --- | --- | --- | --- | --- | --- |
| **Monosaccharides production** | | | | | | | | |
| **Monosaccharide** | **Gene** | | **Protein** | | | **COG ID** | **KEGG ID** | **tblastN results** |
| Glucose | *glgA* | | starch synthase | | | COG0297 | K00703 |  |
|  | *glgC* | | glucose-1-phosphate adenylyltransferase | | | COG0448 | K00975 |  |
| Xylose | *uxs* | | UDP-glucuronate decarboxylase | | | COG0451 | K08678 |  |
| Rhamnose | *rfbA* | | dTDP-glucose pyrophosphorylase | | | COG1209 | K00973 |  |
|  | *rfbB* | | dTDP-D-glucose 4,6-dehydratase | | | COG1088 | K01710 |  |
|  | *rfbC* | | dTDP-4-dehydrorhamnose 3,5-epimerase | | | COG1898 | K01790 |  |
|  | *rfbD* | | dTDP-4-dehydrorhamnose reductase | | | COG1091 | K00067 |  |
| Arabinose | *uxe* | | UDP-arabinose 4-epimerase | | |  |  | EKU99180.1  Query cover: 92%  Identity: 71% |
| Fucose | *fcl* | | GDP-L-fucose synthase | | | COG0451 | K02377 |  |
| Mannose | *manA* | | mannose-6-phosphate isomerase | | | COG0662 | K01809 |  |
| Galactose | *galE* | | UDP-glucose 4-epimerase | | | COG1087 | K01784 |  |
| Fructose | *pgi* | | Glucose-6-phosphate isomerase | | | COG0166 | K01810 |  |
| Ribose | *rpiA* | | ribose 5-phosphate isomerase A | | | COG0120 | K01807 |  |
| Galactosamine | *wbpP* | | UDP-N-acetylgalactosamine 4-epimerase | | |  |  | OWY73298.1  Query cover: 100%  Identity: 37.38% |
| Glucosamine | *glmU* | | glucosamine-1-phosphate N-acetyltransferase | | | COG1207 | K04042 |  |
| Galacturonic acid | *cap1J* | | UDP-glucuronate 4-epimerase | | |  |  | EHA62317.1  Query cover: 100%  Identity: 35.91% |
| Glucuronic acid | *ugd* | | UDP glucose 6-dehydrogenase | | | COG1004 | K00012 |  |
| **EPS assembly and export: Wzy dependent pathway** | | | | | | | | |
| **Protein** | **Gene** | | **Function** | | | **COG ID** | **KEGG ID** | **tblastN results** |
| Outer membrane  polysaccharide export (OPX) protein | *wza* | | Translocation across the outer membrane | | | COG1596 | K01991 |  |
| Phosphatase | *wzb* | | Polymerization and translocation | | | COG0394 | K25307 |  |
| Polysaccharide copolymerase (PCP) | *wzc* | | Polymerization and translocation | | | COG3206 | K16554 |  |
| Polysaccharide flippase | *wzx* | | Flips nucleotide units across the membrane | | |  |  | WP_011153586.1  Query cover: 98%  Identity: 53.65% |
| Polysaccharide polymerase | *wzy* | | Assembly of the polysaccharide chain | | |  |  | WP_010874002.1 Query cover: 95%  Identity: 37.29% |
| **EPS assembly and export:**  **ABC-transporter dependent pathway** | | | | | | | | |
| **Protein** | **Gene** | | **Function** | | | **COG ID** | **KEGG ID** | **tblastN results** |
| ATP-binding protein (ABC transporter) | *kpsT* | | Transfer of polysaccharides across the cytoplasmic membrane | | | COG1134 | K09691 |  |
| Transport permease protein (ABC transporter) | *kpsM* | | Transfer of polysaccharides across the cytoplasmic membrane | | | COG1682 | K09690 |  |
| Polysaccharide copolymerase (PCP) | *kpsE* | | Translocation across the outer membrane and polymerization | | |  |  | BAA10474.1  Query cover: 89%  Identity: 73% |
| Outer membrane  polysaccharide export (OPX) | *kpsD* | | Translocation across the outer membrane | | |  |  | AVZ30903.1  Query cover: 96%  Identity: 46.79% |
| **EPS assembly and export: synthase dependent pathway** | | | | | | | | |
| **Protein** | **Gene** | | **Function** | | | **COG ID** | **KEGG ID** | **tblastN results** |
| Polymerase/synthase | *alg8* | | Synthesis, polymerization and export across the plasma membrane | | | COG2274 | K11004 |  |
| Mannuronan synthase | *alg44* | | Regulates alg8 | | | COG0845 | K11003 |  |
| **Photoprotection** | | | | | | | | |
| **Carotenoids production** | | | | | | | | |
| **Protein** | | **Gene** | | **Function** | **COG ID** | | **KEGG ID** | **tblastN results** |
| geranylgeranylpyrophosphate (GGPP) synthase | | *crtE* | | Synthesis of GGPP | COG0142 | | K13789 |  |
| phytoene synthase | | *crtB* | | Synthesis of phytoene | COG1562 | | K02291 |  |
| phytoene desaturase | | *crtP* | | Synthesis of phytofluene | COG3349 | | K02293 |  |
| zeta‐carotene desaturase | | *crtQ* | | Synthesis of neurosporene | COG3349 | | K00514 |  |
| prolycopene isomerase | | *crtH* | | Synthesis of lycopene | COG1233 | | K09835 |  |
| lycopene cyclase | | *cruA* | | Synthesis of β-carotene | COG0644 | | K14605 |  |
| β-carotene hydroxylase | | *crtR* | | Synthesis of zeaxanthin | COG3239 | | K02294 |  |
| β-carotene ketolase | | *crtW* | | Synthesis of echinenone | COG3239 | | K09836 |  |
| β-carotene ketolase | | *crtO* | | Synthesis of canthaxanthin | COG1233 | | K02292 |  |
| **Scytonemin production** | | | | | | | | |
| **Scytonemin production** | | **Gene** | | **Function** | **COG ID** | | **KEGG ID** | **tblastN results** |
| Prephenate dehydrogenase | | *tyrA* | | Phephenate dehydrogenase | COG0287 | | K14187 |  |
| Scytonemin biosynthesis protein | | *scyA* | | Synthesis of a β-ketoacid compound |  | |  | WP_012408018.1  Query cover: 86%  Identity: 25.45% |
| Tryptophan dehydrogenase | | *scyB* | | Synthesis of indole-3-pyruvic acid |  | |  | QHG19687.1  Query cover: 75%  Identity:27.83% |
| ScyD/ScyE family protein | | *scyD/E* | | Oxidative dimerization |  | |  | WP_012408014.1  Query cover: 28%  Identity:25.56% |
| 3-dehydroquinate synthase | | *aroB* | | Start of shikimic acid pathway | COG0337 | | K01735 |  |
| DAHP synthase | | *aroG* | | Start of shikimic acid pathway | COG2876 | | K03856 |  |
| Several proteins | | *trpA*  *trpB*  *trpC*  *trpD*  *trpE*  *trpF*  *trpG* | | Involved in the tryptophan biosynthesis | COG0159  COG0133  COG0134  COG0547  COG0147  COG0135  COG0512 | | K01695  K01696  K01609  K00766  K01657  K01817  K01658 |  |
| **Mycosporine-like aa production** | | | | | | | | |
| **Protein** | | **Gene** | | **Function** | **COG ID** | | **KEGG ID** | **tblastN results** |
| 2-demethyl-4-deoxygadusol synthase | | *mysA* | | Synthesis of DDG (2-demethyl-4-deoxigadusol) |  | |  | Q3M6C3.1  Query cover: 73%  Identity: 29.3% |
| O-methyltransferase | | *mysB* | | Synthesis of 4-DG (4-deoxygadusol) |  | |  | BBB38337.1  Query cover: 80%  Identity: 39.13% |
| Non ribosomal peptide synthase | | *mysE* | | Shinorine production |  | |  | ABA23460  Query cover: 98%  Identity:39.61% |
| **Oxidative stress protection** | | | | | | | | |
| **Protein** | | **Gene** | | **Function** | **COG ID** | | **KEGG ID** | **tblastN results** |
| Orange carotenoid-binding protein | | *ocp1* | | Nonphotochemical quenching and thermal dissipation of high light energy |  | |  | NEQ30519.1 Query cover: 100%  Identity:78.06% |
| D1 protein of PSII | | *psbA* | | Oxidative stress acclimation |  | | K02703 |  |
| Superoxide dismutase Fe-Mn | | *sod* | | Antioxidant enzyme | COG0605 | | K04564 |  |
| DNA- binding ferritin-like protein | | *dps* | | Protects DNA from oxidative damage | COG0783 | | K04047 |  |
| Thioredoxin | | *trxA* | | Regulates GroEL | COG3118 | | K03671 |  |
| Methionine sulfoxide reductase | | *msrA* | | Reverse protein oxidation | COG0225 | | K07304 |  |
| Mn-containing catalase | | *cat* | | ROS scavenger | COG3546 | | K07217 |  |
| Peroxiredoxin | | *prx* | | Oxidative stress protection | COG1225 | | K03386 |  |
| Chaperone | | *dnaK* | | Proteins folding | COG0443 | | K04043 |  |
| Chaperone | | *dnaJ* | | Proteins folding | COG0484 | | K05516 |  |
| Chaperonin | | *groEL* | | Proteins folding | COG0459 | | K04077 |  |
| Chaperonin | | *groES* | | Proteins folding | COG0234 | | K04078 |  |
| Chaperon | | *htpG* | | Proteins folding | COG0326 | | K04079 |  |
| ATP-dependent chaperone | | *clpB* | | Proteins folding | COG0542 | | K03695 |  |
| Histidine kinase | | *hik34* | | Regulates heat shock proteins | COG0642 | |  |  |
| **DNA repair** | | | | | | | | |
| **Protein** | | **Gene** | | **Function** | **COG ID** | | **KEGG ID** | **tblastN results** |
| DNA polymerase III | | *holB* | | Homologous recombination and MMR pathways | COG0470 | | K02341 |  |
| ATP-dependent DNA helicase RecQ | | *recQ* | | Homologous recombination repair pathway | COG0514 | | K03654 |  |
| ATP-dependent DNA helicase RecG | | *recG* | | Homologous recombination repair pathway | COG1200 | | K03655 |  |
| Single-stranded-DNA-specific exonuclease | | *recJ* | | Homologous recombination and MMR pathways | COG0608 | | K07462 |  |
| Single-strand DNA-binding protein | | *ssb* | | Homologous recombination and MMR pathways | COG0629 | | K03111 |  |
| Recombinational DNA repair proteins | | *recORF* | | Homologous recombination repair pathway | COG1381  COG1195  COG0353 | | K03584  K03629  K06187 |  |
| Recombinase | | *recA* | | Homologous recombination repair pathway | COG0468 | | K03553 |  |
| Hollyday junction enzymes | | *uvrABC* | | Homologous recombination and NER repair pathways | COG0632  COG2255  COG0817 | | K03550  K03551  K01159 |  |
| Primosomal protein | | *priA* | | Homologous recombination repair pathway | COG1198 | | K04066 |  |
| Photolyase | | *pL* | | PL repair mechanism | COG1533 | | K03716 |  |
| Nucleotide glycosylase | | *fpg* | | BER pathway | COG0266 | | K10563 |  |
| Endonuclease VIII | | *nei* | | BER pathway | COG0266 | | K05522 |  |
| Endonuclease III | | *nth* | | BER pathway | COG0177 | | K10773 |  |
| DNA polymerase I | | *dpoI* | | BER and NER pathways | COG0749 | | K02335 |  |
| DNA ligase | | *lig* | | BER, NER and MMR pathways | COG0272 | | K01972 |  |
| Exonuclease III | | *xth* | | BER pathway | COG0708 | | K01142 |  |
| DNA-3-methyladenine glycosylase | | *mpg* | | BER pathway | COG2094 | | K03652 |  |
| DNA-3-methyladenine glycosylase II | | *alkA* | | BER pathway | COG0122 | | K01247 |  |
| A/G-specific adenine glycosylase | | *mutY* | | BER pathway | COG1194 | | K03575 |  |
| Uracil-DNA glycosylase | | *udg* | | BER pathway | COG1573 | | K21929 |  |
| DNA helicase | | *uvrD* | | NER and MMR pathways | COG0210 | | K03657 |  |
| Transcription-repair coupling factor | | *mfd* | | NER pathway | COG1197 | | K03723 |  |
| DNA mismatch repair ATPase | | *mutS* | | MMR pathway | COG0249 | | K03555 |  |
| DNA mismatch repair ATPase | | *mutL* | | MMR pathway | COG0323 | | K03572 |  |
| Exonuclease VII (large and small subunits) | | *exo VII* | | MMR pathway | COG1570  COG1722 | | K03601  K03602 |  |
| Site-specific DNA-adenine methylase | | *dam* | | MMR pathway | COG0338 | | K06223 |  |
| Transcriptional repressor | | *lexA* | | SOS system | COG1974 | | K01356 |  |
| **Osmotic stress protection** | | | | | | | | |
| **Salt-out strategy** | | | | | | | | |
| **Protein** | | **Gene** | | **Function** | **COG ID** | | **KEGG ID** | **tblastN results** |
| Na+ antiporter | | *nhA* | | Export Na+ outside cells | COG0025 | | K24160 |  |
| K+ transport protein | | *ktrA* | | Import K+ inside cells | COG0569 | | K03499 |  |
| K+ transport protein | | *ktrB* | | Import K+ inside cells | COG0168 | | K03498 |  |
| Glycosyltransferase | | *ktrE* | | Import K+ inside cells |  | |  | BAA18015.1  Query cover: 98%  Identity: 61.64% |
| K+ sensing histidine kinase | | *kdpD* | | Activates *kdp* genes expression | COG2205 | |  |  |
| **Osmoprotectants production and uptake** | | | | | | | | |
| **Protein** | | **Gene** | | **Function** | **COG ID** | | **KEGG ID** | **tblastN results** |
| Sucrose-6-phosphatase | | *spp* | | Sucrose synthesis | COG0561 | | K07024 |  |
| Sucrose-P-synthase | | *sps* | | Sucrose synthesis |  | |  | WP_044521731.1 Query cover: 97%  Identity: 67.72% |
| ABC-type sugar transport system | | *ggtA* | | Sucrose/Trehalose/  Glucosylglycerol uptake | COG3839 | | K10112 |  |
| maltooligosyl trehalose synthase | | *treY* | | Trehalose synthesis | COG3280 | | K06044 |  |
| maltooligosyl trehalose trehalohydrolase | | *treZ* | | Trehalose synthesis | COG0296 | | K01236 |  |
| isoamylase | | *treX* | | Trehalose synthesis | COG1523 | | K01214 |  |
| starch synthase | | *glgA* | | Trehalose synthesis | COG0297 | | K00703 |  |
| 1,4-alpha-glucan branching enzyme | | *glgB* | | Trehalose synthesis | COG0296 | | K00700 |  |
| glucose-1-phosphate adenylyltransferase | | *glgC* | | Trehalose synthesis | COG0448 | | K00975 |  |
| sorbitol-6-phosphate 2-dehydrogenase | | *srlD* | | Sorbitol synthesis |  | |  | NP_417185.1  Query cover: 99%  Identity: 32.56% |
| Glutamate synthase | | *gltS* | | Glutamate synthesis | COG0067 | | K00284 |  |
| Glutamine-binding periplasmic protein | | *glnH* | | Glutamate/glutamine ABC transporter |  | |  | WP_010997579.1  Query cover: 92%  Identity: 54.98% |
| Glutamine-binding periplasmic protein | | *glnP* | | Glutamate/glutamine ABC transporter |  | |  | WP_010997338.1  Query cover: 45%  Identity: 27.31 % |
| ATP-binding protein | | *glnQ* | | Glutamate/glutamine ABC transporter |  | |  | GAP98062.1  Query cover: 99%  Identity: 39.37% |
| ATP-binding protein | | *gluA* | | Glutamate ABC transporter |  | |  | EGY1326349.1  Query cover: 100%  Identity: 39.83% |
| N-I type | | *natA*  *natB*  *natC*  *natD*  *natE* | | Aminoacid transporter | COG0411  COG0683  COG4177  COG0559  COG0410 | | K11957  K11954  K11955  K11956  K11958 |  |
| N-II type | | *natF* | | Aminoacid transporter |  | |  | BAB75863.1  Query cover: 57%  Identity: 26.99% |
| glutamate-5-semialdehyde dehydrogenase | | *proA* | | Proline synthesis | COG0014 | | K00147 |  |
| glutamate 5-kinase | | *proB* | | Proline synthesis | COG0263 | | K00931 |  |
| pyrroline-5-carboxylate reductase | | *proC* | | Proline synthesis | COG0345 | | K00286 |  |
| ATP-binding protein | | *opuA* | | Proline/Glycine Betaine transporter | COG1125 | | K05847 |  |
| permease protein | | *opuB* | | Proline/Glycine Betaine transporter | COG1174 | | K05846 |  |
| substrate-binding protein | | *opuC* | | Proline/Glycine Betaine transporter | COG1732 | | K05845 |  |
| Choline dehydrogenase | | *betA* | | Glycine betaine synthesis (choline pathway) | COG2303 | |  |  |
| NAD-dependent aldehyde dehydrogenase | | *betB* | | Glycine betaine synthesis (choline pathway) | COG1012 | |  |  |
| glycine N-methyltransferase | | *GNMT* | | Glycine betaine synthesis (glycine pathway) |  | |  | BAS66417.1  Query cover: 60%  Identity: 36.04% |
| dimethylglycine N-methyltransferase | | *DMT* | | Glycine betaine synthesis (glycine pathway) |  | |  | WP_015227494.1  Query cover: 64%  Identity: 32.22% |
| L-proline glycine betaine binding ABC transporter | | *proX* | | Ectoine/glycine betaine/proline/choline uptake |  | |  | GAP95390.1  Query cover: 93%  Identity: 48.35% |
| membrane component | | *proW* | | Ectoine/glycine betaine/proline/choline uptake |  | |  | TVP68275.1  Query cover: 100%  Identity: 30.19% |
| L-proline glycine betaine ABC transport system permease | | *proV* | | Ectoine/glycine betaine/proline/choline uptake |  | |  | GAP95388.1  Query cover: 100%  Identity: 61.09% |
